# Applying gradient tree boosting to QTL mapping with Shapley additive explanations

**DOI:** 10.1101/2024.01.15.575690

**Authors:** Tomohiro Ishibashi, Akio Onogi

## Abstract

Mapping quantitative trait loci (QTLs) is one of the major goals of quantitative genetics; however, identifying the interactions between QTLs (i.e., epistasis) remains challenging. Recently developed machine learning methods, such as deep learning and gradient boosting, are transforming the real world. These methods could advance QTL mapping methodologies because of their high capability for capturing complex relationships among features. One problem with applying such complex models to QTL mapping is the evaluation of feature importance. In this study, XGBoost, a popular gradient tree boosting algorithm, was applied for QTL mapping in biparental populations with Shapley additive explanations (SHAPs). SHAP is a local (i.e., instance-wise) importance index with the desired properties as feature importance indices. The SHAP-assisted XGBoost (SHAP-XGB) was compared with conventional methods, including composite interval mapping (CIM), multiple interval mapping (MIM), inclusive CIM (ICIM), and BayesC, using simulations and rice heading date data. SHAP-XGB performed comparablely to CIM, MIM, ICIM, and BayesC in mapping main QTL effects and was superior to MIM, ICIM, and BayesC in mapping QTL interaction effects. As SHAP can evaluate local importance, interactions between markers can be visualized by plotting SHAP interaction values for each instance (plant/line). These results illustrated the strength of SHAP-XGB in detecting and interpreting epistatic QTLs and suggest the possibility that SHAP-XGB complements conventional methods.

## Introduction

Detecting quantitative trait loci (QTLs) underpinning variations in quantitative traits is one of the major goals in analyzing quantitative traits. Detecting QTLs can lead to useful applications such as efficient selection based on QTL genotypes, genomic engineering to modify phenotypes, development of medical technologies by understanding molecular mechanisms, and advancing quantitative genetic theories by deciphering the genetic architecture of quantitative traits (Falconer and Mackay 1996). Interval mapping (IM) is a typical method to detect QTLs in inbred line crosses (Lander and Botstein 1989). IM tests associations of each marker and marker interval with the phenotypes independently. In practice, composite IM (CIM), an extension of IM, is the most popular method (Jansen and Stam 1994, Zeng 1994). CIM considers markers linked with major QTLs as cofactors to enhance detection power and precision when multiple QTLs exist.

Interactions between QTLs, known as epistasis, have also been of interest in quantitative genetics because genes (proteins) function in cooperation. Detecting epistatic QTLs will broaden the applications of the QTLs described above and will be helpful in depicting gene networks underlying complex quantitative traits. A successful example of QTL and epistatic QTL mapping in plant breeding is the rice heading date. To date, a number of QTL/genes and their networks under short- and long-day conditions have been elucidated (Hori et al. 2016, Wei et al. 2020). Methods for detecting epistatic QTLs include multiple interval mapping (MIM, Kao et al. 1999, Laurie et al. 2014), inclusive CIM (ICIM, Li et al. 2008), and Bayesian regression where whole genome-wide markers are used as explanatory variables simultaneously (Zhang and Xu 2005, Li and Sillanpää 2012). A method to detect interactions between QTLs and genetic background was also proposed (Jannink and Jansen 2001). Detection of QTL/gene interactions has been of interest in genome-wide association studies (GWAS), although detection is more difficult than in inbred line crosses because of the generally low allele frequencies. These methods include hierarchical Bayesian modeling for high-order gene interactions (Bayesian high-order interaction toolkit (BHIT) (Wang et al. 2015) and a marginal epistasis test (MAPIT) (Crawford et al. 2017).

Recent machine learning models, such as gradient boosting and deep neural networks, have shown high predictive abilities in various applications. Because these models are inherently capable of capturing complex interactions among features, they are expected to be useful for QTL mapping, particularly for complex QTL interactions. However, unlike linear regressions, the contributions of each feature and feature interaction are not explicitly output. Thus, such complex models should be used with efficient and reliable methods to extract the importance of features for selection. For example, Johnsen et al. (2021) applied Shapley additive explanations (SHAP) to estimate the feature importance of extreme gradient boosting (XGBoost) (Chen and Guestrin 2016) in a GWAS using UK Biobank data. Cui et al. (2022) applied a Shapley-based interaction score (Sundararajan et al. 2020) to deep neural networks to identify gene-gene interactions in a GWAS. The Shapley value was originally proposed to evaluate players’ contributions to rewards in cooperative games (Shapley 1953). By considering players and rewards as features and prediction outcomes, respectively, the Shapley value can be used to evaluate the feature and feature combination importance of the prediction models. SHAP is based on the Shapley value and is characterized by the local importance of the feature; that is, the importance of the feature for each instance (i.e., sample) is estimated (Lundberg and Lee 2017). Because the summation of the SHAP values of the features for an instance is equal to the predicted values of the instance (a property called local accuracy), it is feasible to determine which features have a large impact on the prediction of the instance (Lundberg and Lee 2017, Lundberg et al. 2018). SHAP also has two additional properties–missingness and consistency–that are desirable as feature importance indices.

It may be possible to map QTLs and QTL interactions using SHAP or Shapley value-based scores and recently advanced machine learning more efficiently and effectively than conventional methods. The GWASs mentioned above partially prove this possibility; however, evaluation studies are limited. Therefore, several issues remain unresolved. For example, the effect of overfitting complex models to data on the efficiency of QTL mapping remains unclear. Moreover, the translation of SHAP values into signals for QTL detection has not been thoroughly examined. The properties of mapping dominance effects are also not examined well. In addition, comparisons with conventional methods such as CIM, MIM, ICIM, and Bayesian regressions have not been conducted. In this study, we evaluated the potential of XGBoost for QTL and epistatic QTL mapping using SHAP. XGBoost is an algorithm for gradient boosting, which is a class of ensemble learning (Chen and Guestrin 2016). XGBoost was selected because it is one of the most prevalent gradient boosting algorithms. We characterized this SHAP-assisted XGBoost (SHAP-XGB) by applying the method to simulated and real biparental populations and comparing it with conventional methods. The results illustrated the strength of SHAP-XGB in detecting and interpreting epistatic QTLs, as expected, and suggested the possibility that SHAP-XGB complements conventional methods.

## Materials and Methods

### Extreme gradient boosting

XGBoost is based on decision trees and represents the response of instance *i* (the phenotype of plant/line *i* in the case of QTL mapping), *y*_*i*_, with the linear combination of *K* decision trees as

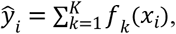

where *f* is a tree with structure *q* and leaf weight *w* and *x*_*i*_ is the vector of feature (marker genotype in the case of QTL mapping). In other words, *y*_*i*_ is given by the sum of the outputs of multiple decision trees. The decision trees are sequentially added at each iteration. Thus, the objective function of XGBoost at iteration *t*, that is, when adding *t*th tree, can be written as

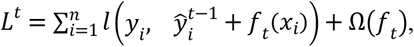

where *n* is the number of instances (plants/lines), *l* is a loss function, 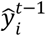 is *ŷ*_*i*_ at iteration *t* − 1, *f*_*t*_ is the tree added at iteration *t*, and Ω is a penalty function on the tree structure (i.e., the number of leaves and their weights). A penalty function is added to avoid overfitting the model. By approximating *l* using Taylor expansions and dropping the constant terms, the objective function becomes

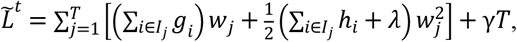

where *g*_*i*_ and *h*_*i*_ are the first and second order gradients of the loss function, *I*_*j*_ is the set of instances assigned to leaf *j, T* is the number of leaves, and *λ* and *γ* are the regularization parameters. The optimal weight of leaf 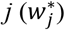 and the corresponding optimal value of 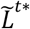 are

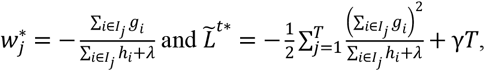

respectively. Using these results, whether a leaf *I* should be split into *I*_*L*_ and *I*_*R*_ can be evaluated with the following equation:

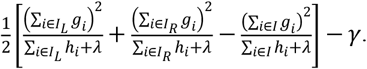

This equation was derived by subtracting 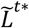 when leaf *I* was split into *I*_*L*_ and *I*_*R*_ from 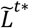 when leaf *I* was a single leaf. Thus, when this equation is positive, 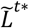 decreases because of splitting. XGBoost uses this formula to search for the tree structures to be added, making the method fast and scalable. The authors proposed exact greedy and approximate algorithms for this type of search.

### Shapley values

In cooperative game theory, the Shapley value measures how much each player contributes to the game’s reward. Assessing the contributions of individual players independently is insufficient in cooperative games because there are usually interactions between players. The core concept of the Shapley value is the ‘marginal contribution’. Marginal contribution represents the extent to which a player’s participation in a cooperative game increases the game’s rewards. This varies according to the combination of players and order of participation. Therefore, the Shapley value of a player is defined by calculating the marginal contribution of a player to all possible player combinations and participation orders and taking their average. In other words, the Shapley value is the average marginal contribution of the player. The Shapley value for player *i* is defined as follows:

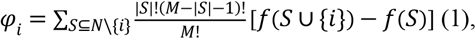

where *N* is a set of players, *S* is a subset of players, *M* is the number of players, and *f* is the function that outputs rewards for the given players.

### SHAP

Players of Shapley values are replaced with features (marker genotypes), and rewards are replaced with the model output (e.g., fitted/predicted phenotypes) in SHAP. The model function to output the rewards given to players takes the form of the expected values, that is, *f*(*S*) in Eq. (1) is expressed as *E*[*f*(*x*)|*x*_*S*_] in SHAP, where *x* and *x*_*S*_ indicate all features and the features in subset *S*, respectively. Thus, the SHAP value of feature *i* of an instance is

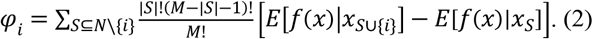

SHAP is an additive feature-contribution method with three properties: local accuracy, missingness, and consistency. Local accuracy indicates that the summation of the feature contributions defined by Eq. (2) is equal to the output of the model. Missingness means that importance is not assigned to a missing feature. Consistency implies that if the model is changed such that a feature has a greater impact on the model, the contribution assigned to that feature does not decrease.

The interaction index was used to assess the impact of one feature of a model when combined with another. The Shapley interaction index between features *i* and *j* is calculated as (Fujimoto et al. 2006),

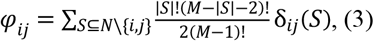

where *δ*_*ij*_(*S*) = *f*(*S* ∪ {*i, j*}) − *f*(*S* ∪ {*i*}) − *f*(*S* ∪ {*j*}) + *f*(*S*). The SHAP interaction value was calculated by replacing the function *f*(*S*) with expectations, as shown in Eq. (2). The SHAP interaction value is split equally between each feature combination, that is, *φ*_*ij*_ = *φ*_*ji*_. Thus, total interaction effects are calculated as *φ*_*ij*_ + *φ*_*ij*_ = 2*φ*_*ij*_. The SHAP value (*φ*_*i*_) of feature *i* for an instance is decomposed to the main effect of the feature (*φ*_*ii*_) and the interaction effects between the feature and the others as

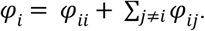

That is, the main effect of feature *i* is obtained by subtracting the SHAP interaction value between feature *i* and *j* (*j* ≠ *i*) from the SHAP value of feature *i*.

Lundberg et al. (2018) proposed an efficient algorithm for calculating SHAP values from tree-based machine learning models (referred to as Tree SHAP). The algorithm exploits the differences in outputs between models with and without a particular feature using a tree structure to evaluate *E*[*f*(*x*)|*x*_*S*_]. The algorithm was implemented in R package xgboost (ver. 1.7.5.1).

### SHAP-XGB algorithm

The SHAP-XGB algorithm is described below and illustrated in Fig. 1. The R scripts for SHAP-XGB are available at https://github.com/Onogi/SHAP-XGB.

**Fig. 1.**
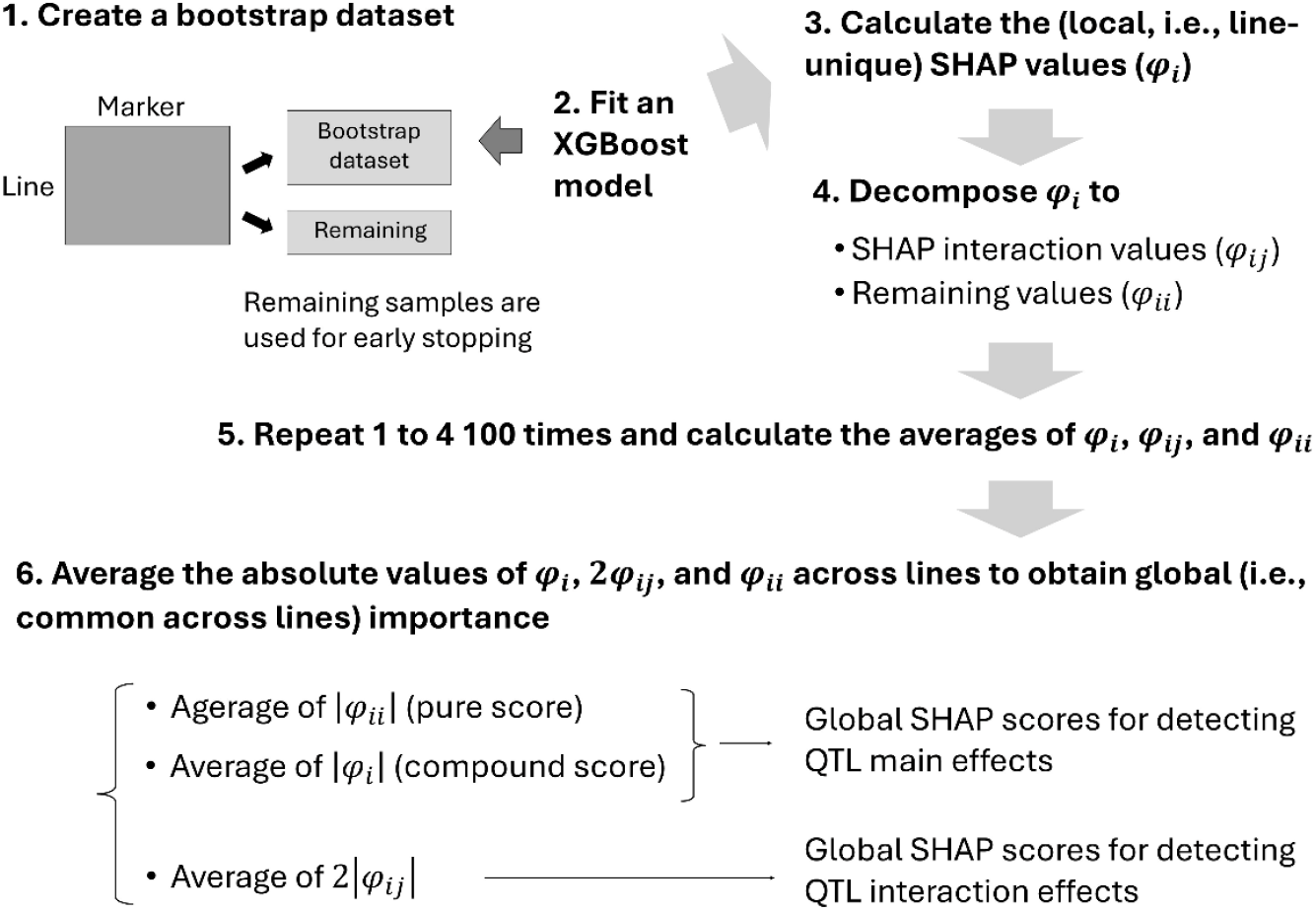
Illustrations of the Shapley additive explanations-assisted XGBoost (SHAP-XGB) algorithm.

**Step 1**: Create a bootstrap dataset.

**Step 2**: Fit an XGBoost model to the dataset, with early stopping determined from the remaining samples. The xgboost R package was used in this study.

**Step 3**: Calculate the SHAP values (*φ*_*i*_) from the fitted samples.

**Step 4**: Decompose the SHAP values into the SHAP interaction values (*φ*_*ij*_) and the remaining values (*φ*_*ii*_).

**Step 5**: Repeat steps 1–4 100 times and calculate the average of *φ*_*i*_, *φ*_*ij*_, and *φ*_*ii*_. Note that these values are measures of local importance (i.e., for each line).

**Step 6**: Average the absolute values of the values obtained in Step 5 across lines to obtain global importance. The averages of the absolute values of *φ*_*i*_ or *φ*_*ii*_ are referred to as the global SHAP scores and the averages of the absolute values of 2*φ*_*ij*_ are referred to as the global SHAP interaction scores.

Five notifications were added to these algorithms. First, although the absolute SHAP values are averaged across lines to obtain the global importance in Step 6, the consistency property still holds for the global importance scores (Lundberg et al. 2018). Second, bootstrapping was conducted to separate the data into training and validation sets, which were used for early stopping to prevent overfitting. Therefore, other resampling methods could be applied. The number of bootstrap replicates (100) was determined arbitrarily. Thus, fewer numbers—even one—may be acceptable, albeit with a loss of reliability of the SHAP values. Third, default values were used for the hyperparameters of xgboost except for the “early_stopping_rounds” argument, which was set to three. This argument specifies the number of rounds in which the boosting iterations stop if the prediction performance does not improve. These settings resulted in a moderate fit of the models to data, as shown later. We also tested settings that lead to underfitting and overfitting and compared them (see the “Algorithm assessment” section). Fourth, there would be two choices for detecting the main effects of QTLs: using SHAP values that contain interaction scores (i.e., *φ*_*i*_) or using SHAP values from which interaction scores are removed (i.e., *φ*_*ii*_). In this study, the former score is referred to as the “compound” score, and the latter is referred to as the “pure” score for detecting QTL main effects. These two scores were compared as described in the “Algorithm assessment” section. Finally, marker genotype probability calculated using recombination rates can be used as feature *x* (explanatory variables) of SHAP-XGB. Thus, interval mapping is possible, which may result in more precise mapping.

### Comparing methods

The SHAP-XGB method was compared with CIM, MIM, ICIM, and BayesC (Habier et al. 2011) using simulated and real data. Note that BayesC and SHAP-XGB used only marker genotypes as covariates and did not use the genotype probabilities of the loci between markers. For the CIM, MIM, and ICIM, the intervals were set to 1 cM.

### Composite interval mapping (CIM)

CIM was performed using the cim function of the R package qtl (ver. 1.60, Broman et al. 2003). The expectation-maximization algorithm was used. The number of covariates was three, and the window size was 10 cM. CIM was used to detect the main QTL effects. We assessed the influence of the number of covariates and window size on the simulated and real data and found that the influence was negligible (data not shown).

### Multiple interval mapping (MIM)

MIM theory was first proposed by Kao et al. (1999). MIM assumes that the genotypic value is the sum of multiple QTLs and QTL interaction effects. QTL genotypes are probabilistically represented using neighboring marker genotypes and recombination frequencies; thus, the MIM model is a finite mixture model. The QTLs and QTL interactions included in the model were searched in a stepwise manner. MIM was conducted using the QTL cartographer (ver. 2.5_011, Wang et al. 2012). We followed the algorithm proposed by Laurie et al. (2014). The algorithm consisted of multiple steps, including a forward search of the main and interaction effects based on Bayesian information criteria, optimization of QTL positions, and testing of QTL effects with a certain significance level (5%).

### Inclusive composite interval mapping (ICIM)

Li et al. (2007) and Li et al. (2008) proposed the ICIM theory. ICIM consists of two steps: in the first step, markers and marker interactions significantly associated with phenotypes are selected with stepwise regression and one- or two-dimensional interval mapping is conducted in the second step by adjusting the phenotypes with the selected markers that are not located in the target intervals. The ICIM was performed using IciMapping (ver. 4.2.53, Meng et al. 2015). The *p*-values of the markers for entering models in stepwise regression (parameter PIN) were set to 0.001 for the main effects (option ICIM-ADD) and 0.0001 for mapping interaction effects (option ICIM-EPI). We attempted a higher PIN value for ICIM-EPI (0.01) in the simulated data; however, the influence was negligible (data not shown).

### BayesC

We used BayesC as a representative Bayesian regression method because it circumvents a drawback relevant to degrees of freedom in parameter estimation (Habier et al. 2011). BayesC analysis was performed using the vigor function of the R package VIGoR (ver. 1.1.0, Onogi and Arakawa 2022). This package uses fast variational Bayesian algorithms to estimate the posterior distributions of Bayesian alphabets. The model included all markers and marker interactions (multiplications of marker genotypes) and estimated their coefficients simultaneously. The prior distribution of a coefficient was normal if the indicator of the marker (or marker combination) was 1 and 0 if the indicator was 0. The prior distributions of indicators were Bernoulli distributions with hyperparameters of 0.05 for main effects and 0.001 for interaction effects. Different prior normal distributions were assumed for main and interaction effects. Bootstrapping was applied to make the inferences about the indicators more robust. The model was fitted to bootstrap samples, and the posterior means of the indicators were averaged across the bootstrap samples. We used 1000 bootstrap samples in this study.

### Simulations

The usefulness of SHAP-XGB was first investigated using simulations that mimicked biparental F_2_ and RIL populations. The conditions of the simulations were as follows: The number of chromosomes was 8, the length of each chromosome was 120 cM, the number of markers (SNPs) per chromosome was 20, the number of QTLs was five, and the number of individuals was 200. Among the 10 QTL combinations, half were assumed to have interaction effects. The main and interaction effects of the QTLs were generated using the standard normal distribution. The broad-sense heritability was set at 0.8. QTL and marker genotypes were generated using the sim.map and sim.cross functions in the R package qtl (ver. 1.60). Markers were assumed to be randomly distributed along with the chromosomes. The QTLs were also assumed to be randomly distributed, except in Scenario 9, as described later. QTL effects and phenotypes were generated using in-house developed R scripts. The number of replications per scenario was 10, which was determined by considering the MIM and ICIM manual operations. When only additive (dominance) effects were simulated, the interaction effects were simulated as additive-by-additive (dominance-by-dominance) effects by multiplying with the corresponding QTL genotypes.

We simulated nine scenarios using these conditions: (1) F_2_ population with only additive effects (QTL genotypes were coded as −1, 0, and 1). (2) RIL population with additive effects only. (3) F_2_ population with only complete dominance effects (QTL genotypes were coded as 0, 1, and 1). (4) F_2_ population with only overdominance effects (QTL genotypes were coded as 0, 1, and 0). (5) F_2_ population with only additive effects and with epistatic variances increased by setting the variance of interaction effects to 16. (6) F_2_ population with only additive effects, but the number of QTLs with nonzero additive effects was limited to two (i.e., three QTLs only had interaction effects without main effects). (7) F_2_ population with only additive effects, of which the size was increased to 400. (8) RIL population with only additive effects, of which the size was increased to 400. For Scenarios 1–8, the QTLs were assigned so as to be one QTL per chromosome and the chromosomes harboring the QTLs were randomly selected. The QTL combinations with interaction effects were randomly selected. In Scenario 9, three QTLs were on the same chromosome with 30 cM intervals, and two QTLs were on a different chromosome with a 30 cM interval. The QTLs within the same chromosome interacted with each other, and one pair of QTLs on different chromosomes also interacted. All of the gene actions were additive. Chromosomes harboring QTLs and QTL positions were randomly determined. Because, as described later, markers within ±10 cM from QTLs were assumed to be positive markers, each QTL island had 10 cM (30 – 10 – 10) intervals that should be regarded as negative. Thus, if the QTLs on the same chromosome are detected as a single QTL, false positives would increase in the intervals. A summary of the simulation scenarios and realized proportions of variances due to the additive and interaction effects are presented in Table 1.

**Table 1.** Simulation scenarios^*a*^.

| Scenario | Pop. | QTL effects<sup>b</sup> |  | Size | Proportion of variance<sup>c</sup> |  | Purpose | Results |
| --- | --- | --- | --- | --- | --- | --- | --- | --- |
|  |  | Main | Interaction |  | V<sub>A</sub> | V<sub>I</sub> |  |  |
| 1 | F<sub>2</sub> | A | A by A | 200 | 0.476 ± 0.104 | 0.328 ± 0.112 | Evaluation of QTL detection power | Figs. 2 and 3 |
|  |  |  |  |  |  |  | Comparison of pure and compound SHAP values | Fig. S1 |
|  |  |  |  |  |  |  | Comparison of XGB settings | Figs. S2 and S3 |
| 2 | RILs | A | A by A | 200 | 0.408 ± 0.175 | 0.385 ± 0.176 | Evaluation of QTL detection power | Figs. 2, 3, and 4 |
|  |  |  |  |  |  |  | Comparison of XGB settings | Figs. S2 and S3 |
| 3 | F<sub>2</sub> | CD | CD by CD | 200 | 0.417 ± 0.222 | 0.407 ± 0.212 | Evaluation of QTL detection power (dominance) | Fig. S4 |
| 4 | F<sub>2</sub> | OD | OD by OD | 200 | 0.462 ± 0.134 | 0.322 ± 0.130 | Evaluation of QTL detection power (dominance) | Fig. S4 |
| 5 | F<sub>2</sub> | A | A by A | 200 | 0.115 ± 0.037 | 0.679 ± 0.032 | Comparison of pure and compound SHAP values | Fig. S1 |
| 6<sup>d</sup> | F<sub>2</sub> | A | A by A | 200 | 0.313 ± 0.178 | 0.476 ± 0.176 | Comparison of pure and compound SHAP values | Fig. S1 |
| 7 | F<sub>2</sub> | A | A by A | 400 | 0.630 ± 0.091 | 0.175 ± 0.087 | Evaluation of QTL detection power in larger populations | Figs. 2 and 3 |
| 8 | RILs | A | A by A | 400 | 0.383 ± 0.218 | 0.423 ± 0.216 | Evaluation of QTL detection power in larger populations | Figs. 2 and 3 |
| 9 | F<sub>2</sub> | A | A by A | 200 | 0.283 ± 0.183 | 0.398 ± 0.126 | Evaluation of detection power for linked QTLs | Fig. S5 |

<sup>a</sup>All populations were assumed to be derived from biparental crosses. SHAP: Shapley additive explanations; XGB: XGBoost.
<sup>b</sup>A: additive, CD: complete dominance, OD: overdominance.
<sup>c</sup>V<sub>A</sub>: additive variance, V<sub>I</sub>: epistasis variance.
<sup>d</sup>Scenario 6 simulated five QTLs, of which three only had interaction effects.

The accuracy of QTL detection was evaluated using receiver operating characteristic (ROC) curves and the area under the curve (AUC). When drawing ROC curves, markers (or marker intervals in the case of interval mapping) located within ±10 cM of the QTLs were regarded as positive markers. For interaction effects, combinations of markers (or marker intervals) located within ±10 cM of QTLs relevant to the interactions were regarded as positive combinations. On average, 3.4 (±2.2) markers were included in the 20 cM windows and 146.9 (±5.0) markers were outside of the windows. ROC curves and AUC were calculated using the R package pROC ver. 1.18.0 (Robin et al. 2011). Differences in the AUC values between the methods were tested using the Wilcoxon signed-rank test (the significance level was 5% with Bonferroni correction). The global SHAP and global SHAP interaction scores were regarded as QTL signals for SHAP-XGB. The LOD scores were regarded as the signals for CIM and ICIM, respectively, and the bootstrap means of the marker indicators were regarded as the signals for BayesC. Because MIM implemented in QTL Cartographer returns a list of detected QTLs and QTL interactions, ROC curves could not be drawn, and consequently, AUC could not be calculated (note that ROC curves require scores of all variables). Instead, the true and false positive rates were calculated.

### Mapping dominance effects

Whereas the cim function of the qtl package, MIM by the QTL cartographer, and ICIM by QTL IciMapping test both the additive and dominance effects automatically, BayesC by vigor and SHAP-XGB need a specification which effect is targeted. Thus, for BayesC and SHAP-XGB, genotype codes were modified depending on the target effects. The genotypes were coded as [−1, 0, 1], [0, 1, 1], and [0, 1, 0] to test the additive, complete dominance, and overdominance effects, respectively.

### Algorithm assessment

The performance of the compound and pure global SHAP scores in detecting QTL main effects was compared using simulation scenarios (1) (F_2_ population with only additive effects), (5) (F_2_ population with only additive effects and with increased epistatic variances), and (6) (F_2_ population with only additive effects, but the number of QTLs that had non-zero additive effects was limited to two).

Two additional settings of XGBoost were compared to investigate the effects of the degree of fit of the XGBoost models to the data on the QTL detection power. In the underfitting setting, the learning rate “eta” was set to 0.1 (default is 0.3) and the “max_depth,” which specifies the maximum depth of trees, was set to 3 (the default is 6). The “early_stopping_rounds” was set to 3. In the overfitting setting, the hyperparameters were the default values and the number of rounds (boosting iterations) was set to 50 without early stopping. Simulation Scenarios 1 and 2 were used for comparison.

### Real data

The real data were obtained from Onogi et al. (2016). Briefly, days to heading of rice were evaluated for 174 Koshihikari/Kasalath//Koshihikari backcross inbred lines (BILs) and their parental lines. The experiments were conducted in six experimental fields in Japan and Vietnam over three years, resulting in nine environments. The nine environments are summarized in Table 2, and details were provided in Table S1 in Onogi et al. (2016). The lines were genotyped for 162 biallelic markers. The real datasets are shared at https://github.com/Onogi/HeadingDatePrediction.

**Table 2.** Nine environments in the rice heading date data^*a*^.

| Environment name | Location | Year | Transplanting date |
| --- | --- | --- | --- |
| Tsukuba2007 | Tsukuba (N36°01' /E140°06') | 2007 | 8 Jun. |
| Fukuoka2008 | Fukuoka (N33°37' /E130°28') | 2008 | 24 Jun. |
| HaNoi2008 | HaNoi (N21°01' /E105°80') | 2008 | 3 Jul. |
| Ishigaki2008 | Ishigaki (N24°23' /E124°11') | 2008 | 17 Jul. |
| Ishikawa2008 | Ishikawa (N36°30' /E136°36') | 2008 | 16 May |
| ThaiNguyn2008 | ThaiNguyn (N21°35' /E105°49') | 2008 | 15 Jul. |
| Tsukuba2008E | Tsukuba (N36°01' /E140°06') | 2008 (Early sowing) | 25 Apr. |
| Tsukuba2008L | Tsukuba (N36°01' /E140°06') | 2008 (Late sowing) | 9 Jul. |
| Tsukuba2009 | Tsukuba (N36°01' /E140°06') | 2009 | 21 May |

<sup>a</sup>Details of these environments are provided in Table S1 of Onogi et al. (2016).

### Dependency plots

SHAP interaction values were calculated for each instance (line) because SHAP was originally proposed as a local importance measure. Thus, the interactions between QTLs (markers) can be visualized by plotting the SHAP interaction values of the marker combinations of interest for all lines. These plots are referred to as dependency plots. Dependency plots were constructed using the R package SHAPforxgBoost (ver. 0.1.1) with some modifications.

## Results

There are two choices when applying SHAP values for mapping QTL main effects: using compound global SHAP scores that contain interaction scores or using pure global SHAP scores from which interaction scores are removed. We first compared these two types of values for mapping the QTL main effects in simulation Scenarios 1, 5, and 6. Scenario 1 simulated the F_2_ populations with additive and additive-by-additive interactions. Scenarios 1 and 5 differed in the proportion of phenotypic variance explained by epistatic variance (0.328 vs. 0.679 on average; Table 1), and Scenarios 1 and 6 differed in the number of QTLs that had main effects (5 vs. 2; Table 1). In Scenario 6, three QTLs only had interaction effects. The AUC values and ROC curves were comparable between the compound and pure scores (Supplemental Fig. 1). A slight increase in the detection power was observed for the compound scores in Scenario 5. In contrast, the false positive rates increased slightly for the compound scores in Scenario 6. This is reasonable because the three QTLs lacked main effects and only had interaction effects in Scenario 6. Thus, we concluded that the compound SHAP scores were comparable with or even better than the pure SHAP scores in QTL mapping as long as the QTLs that had non-zero main effects were involved in the interactions. Hereafter, we used compound scores to map the main QTL effects.

Next, we investigated the effect of the fitting degree of the XGBoost models on QTL mapping accuracy. We compared the three settings that led to underfitting, moderate fitting, and overfitting in simulation Scenarios 1 and 2. The SHAP values were either underestimated when underfitting was induced and overestimated when overfitting was induced (Supplemental Fig. 2). Nevertheless, as the relative values between the markers did not vary significantly, the AUC values and ROC curves were hardly changed by the fitting status (Supplemental Fig. 3). We used a setting that induced a moderate fit.

In comparative analyses mapping main QTL effects, SHAP-XGB generally showed higher or comparable AUC values than CIM, ICIM, and BayesC (Fig. 2). When compared with MIM, SHAP-XGB showed true positive rates comparable to MIM at the false positive rates that MIM showed (Fig. 2). In comparative analyses mapping QTL interaction effects, SHAP-XGB showed significantly higher AUC values than ICIM and BayesC (Fig. 3). When compared with MIM, SHAP-XGB showed higher true positive rates than MIM under false positive rates comparable to those of MIM (Fig. 3), suggesting that SHAP-XGB can detect more QTL interactions than MIM under comparable false positive rates. Interactions between markers (QTLs) can be interpreted using dependency plots. The SHAP interaction values (local importance metrics) for a replication of Scenario 2 are shown in Fig. 4. The effects of two QTLs, *QTL2* and *QTL3* tagged with *D3M19* and *D4M11* markers, respectively, were inverted depending on the genotype of *QTL4* tagged with *D6M1*. These figures illustrate that dependency plots using local importance scores can facilitate the interpretation of interactions.

**Fig. 2.**
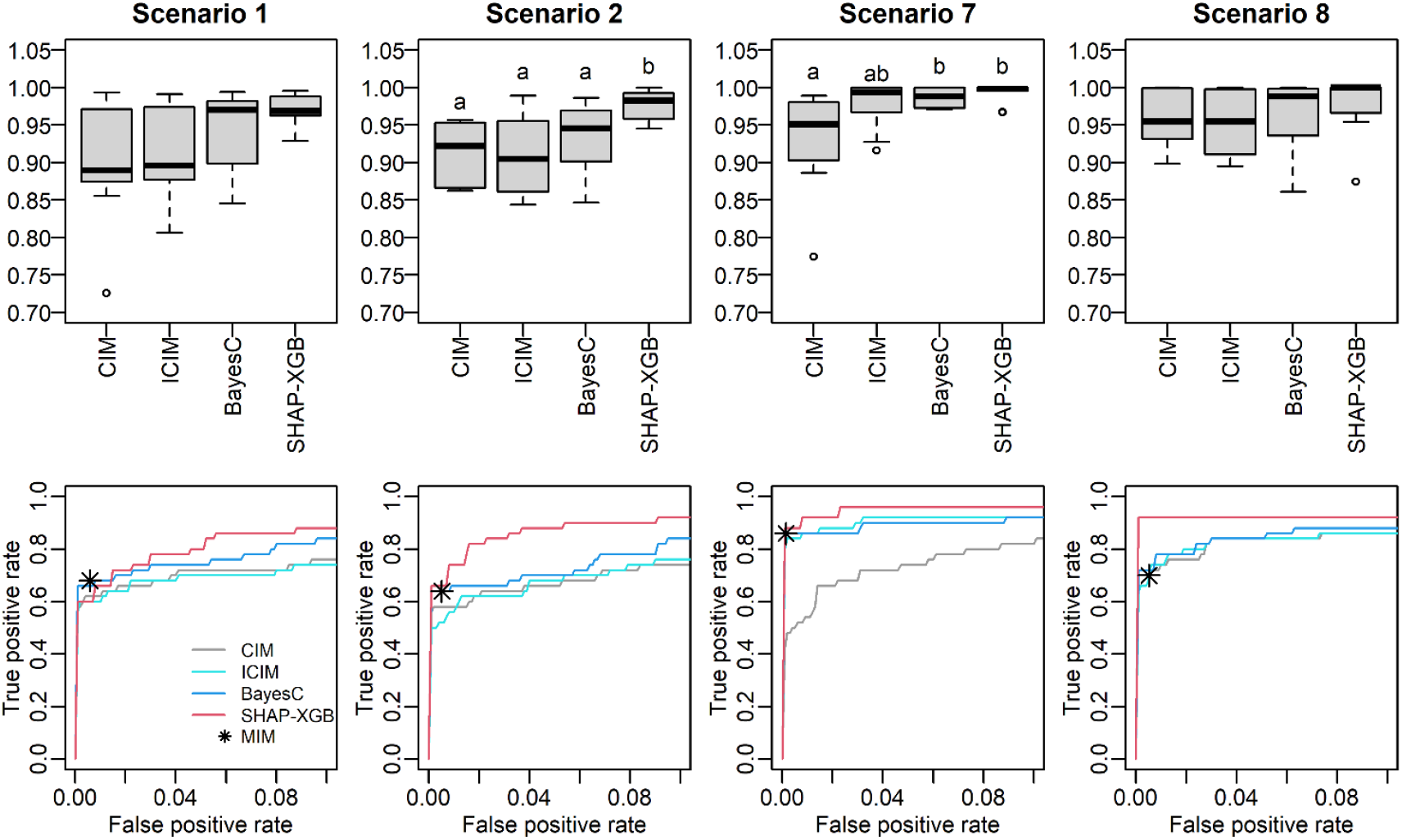
Area under the curve (AUC) values and receiver operating characteristic (ROC) curves in mapping main QTL effects. The top and bottom panels are the AUC values and ROC curves, respectively. The ROC curves are the mean curves of 10 replications. Scenarios 1 and 7 simulated F_2_ populations, and Scenarios 2 and 8 simulated recombinant inbred line populations. The population sizes were 200 for Scenarios 1 and 2, and 400 for Scenarios 7 and 8. Different letters in the box plots denote significant differences (*P* <0.05). Because the results of MIM are the lists of detected QTLs, true and false positives are shown with the asterisk in the bottom plots. CIM: composite interval mapping, ICIM: inclusive CIM, SHAP-XGB: Shapley additive explanations-assisted XGBoost, MIM: multiple interval mapping.

**Fig. 3.**
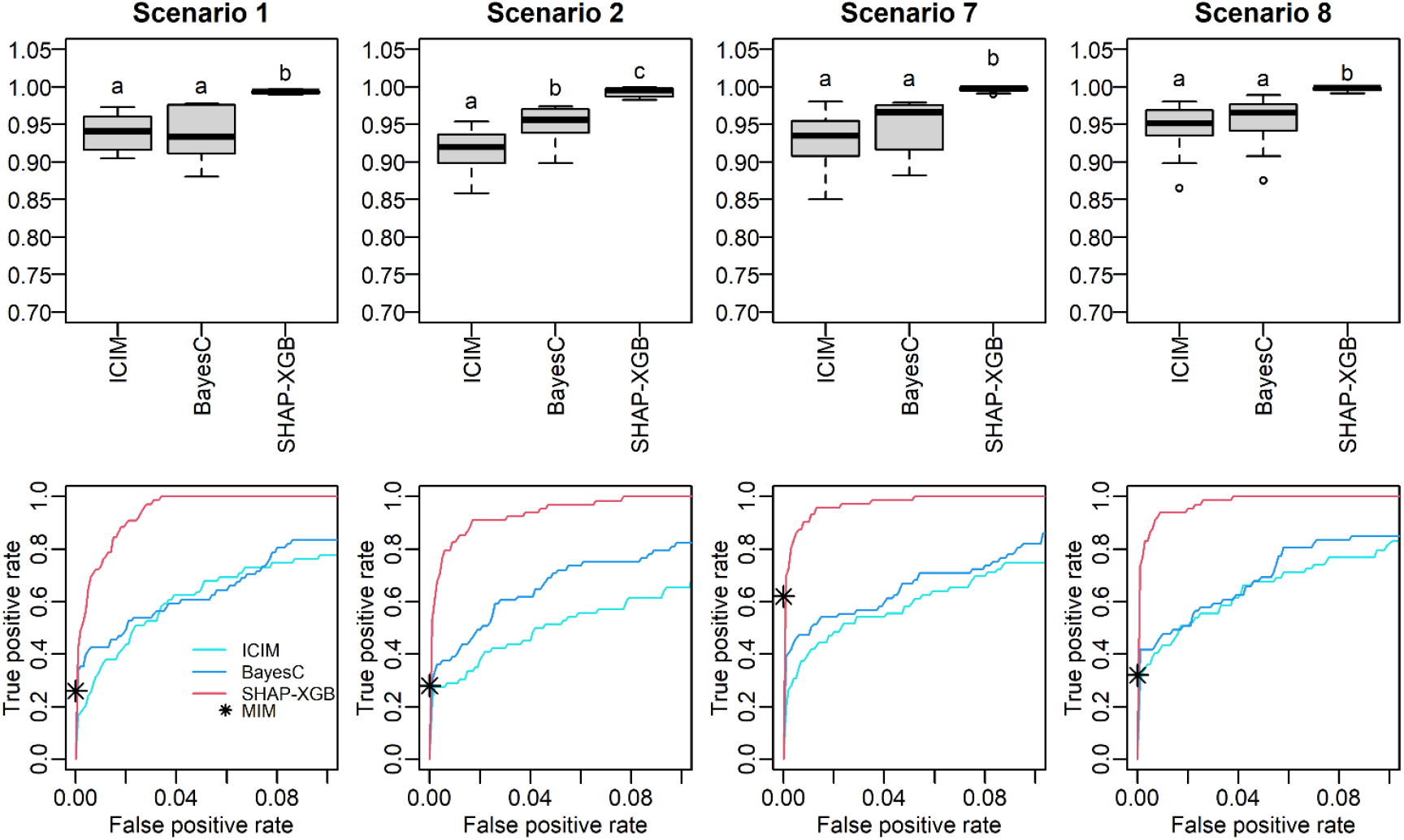
Area under the curve (AUC) values and receiver operating characteristic (ROC) curves in mapping QTL interaction effects. The top and bottom panels are the AUC values and ROC curves, respectively. The ROC curves are the mean curves of 10 replications. Scenarios 1 and 7 simulated F_2_ populations and Scenarios 2 and 8 simulated recombinant inbred line populations. The population sizes were 200 for Scenarios 1 and 2, and 400 for Scenarios 7 and 8. Different letters in the box plots denote significant differences (*P* <0.05). Because the results of MIM are the lists of detected interactions, true and false positives are shown with asterisks in the bottom plots. ICIM: inclusive composite interval mapping, SHAP-XGB: Shapley additive explanations-assisted XGBoost, MIM: multiple interval mapping.

**Fig. 4.**
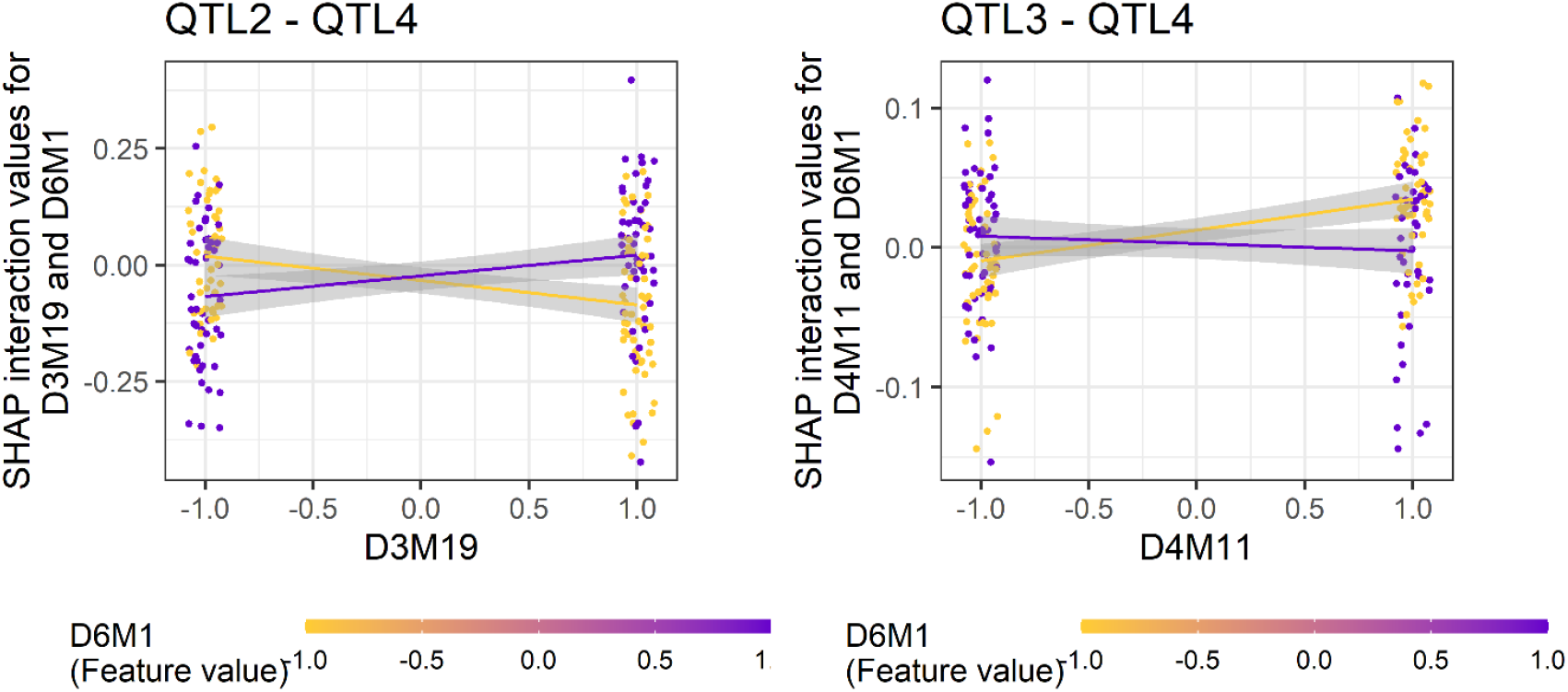
Dependencies between QTLs in a simulated dataset. The Shapley additive explanations (SHAP) interaction values of each line are shown as dots. The left panel shows the interaction between the 2nd and 4th QTLs, and the right panel shows that between the 3rd and 4th QTLs. *D3M19, D4M11*, and *D6M1* are the markers linked to the 2nd, 3rd, and 4th QTLs, respectively. Genotypes of *D6M1* are shown using different colors. Dots are scattered towards the x-axis. The regression lines were added for each genotype of *D6M1*. The shades denote the standard errors. The left panel shows that, when the genotype of *D6M1* is −1, SHAP interaction values (i.e., phenotypes) tend to decrease as the genotype of *D3M19* changes from −1 to 1, and when the genotype of *D6M1* is 1, SHAP interaction values tend to increase. The right panel shows inverted tendencies between *D4M11* and *D6M1*. The gray areas indicate the 95% confidence intervals of regression lines.

SHAP-XGB showed a performance comparable with that of CIM, MIM, ICIM, and BayesC for main effect mapping, and better than that of MIM, ICIM, and BayesC for interaction effect mapping in mapping dominance effects (Scenarios 3 and 4), as observed in mapping additive effects (Supplemental Fig. 4). In addition, SHAP-XGB showed comparable or only slightly decreased AUC values when additive effects were targeted, compared to those when dominance effects were targeted (Supplemental Fig. 4). These results are in contrast to those of BayesC, where misspecification of the target effects resulted in a more prominent decrease in accuracy. These results suggest that SHAP-XGB was insensitive to the effect type. This characteristic was also observed when dominance-by-dominance interactions were targeted. In other words, dominance-by-dominance effects can be detected even when additive-by-additive effects were targeted.

When QTLs are linked to each other, they can be detected as a single QTL if multiple QTL signals cannot be separated. This phenomenon has motivated the development of methods, such as CIM and ICIM. In Scenario 9, where the QTLs were linked, SHAP-XGB showed a comparable performance with CIM and ICIM in mapping the main effects and a slightly better performance than ICIM in mapping the interaction effects (Supplemental Fig 5). Thus, it is suggested that SHAP-XGB can separate signals from linked QTLs, as well as CIM and ICIM.

The XGBoost models explained 94.3% (± 1.9%) of the phenotypic variance with the setting of moderate fitting in the real rice data analyses. SHAP-XGB and the compared methods showed similar results in mapping the main QTL effects (Supplemental Fig. 6 and Fig. 5), and markers linked to *Hd6* (chr. 3, Takahashi et al. 2001), *Hd1* (chr. 6, Yano et al. 1997), and *DTH8* (chr. 8, Wei et al. 2010) showed higher signals for each method. The ICIM results were generally lenient when mapping epistatic QTLs (Supplemental Fig. 7 and Fig. 6), whereas the BayesC results were stringent (or conservative). SHAP-XGB showed intermediate results between ICIM and BayesC, and could pinpoint interacting combinations between known QTLs within reasonably narrow margins. The results of the MIM in the mapping interactions were relatively stringent; nine interactions were detected in nine environments (Supplemental Table 1). In general, these interactions were detected using SHAP-XGB. The global SHAP interaction scores in each environment showed interesting trends across latitudes (Fig. 7). Interactions between *Hd1* and *Hd6, Hd1* and *DTH8*, and *Hd1* and *Hd2*/*OsPRR37* (Yano et al. 1997, Koo et al. 2013) showed higher scores at higher latitudes (environments Tsukuba2007, Tsukuba2008E, Tsukuba2009, Ishikawa2008, and Fukuoka2008), but lower scores at lower latitudes (environments Ishigaki2008, HaNoi2008, and ThaiNguyn2008). In contrast, the interaction between *DTH8* and *Hd2* showed larger scores at lower latitudes but lower scores at higher latitudes.

**Fig. 5.**
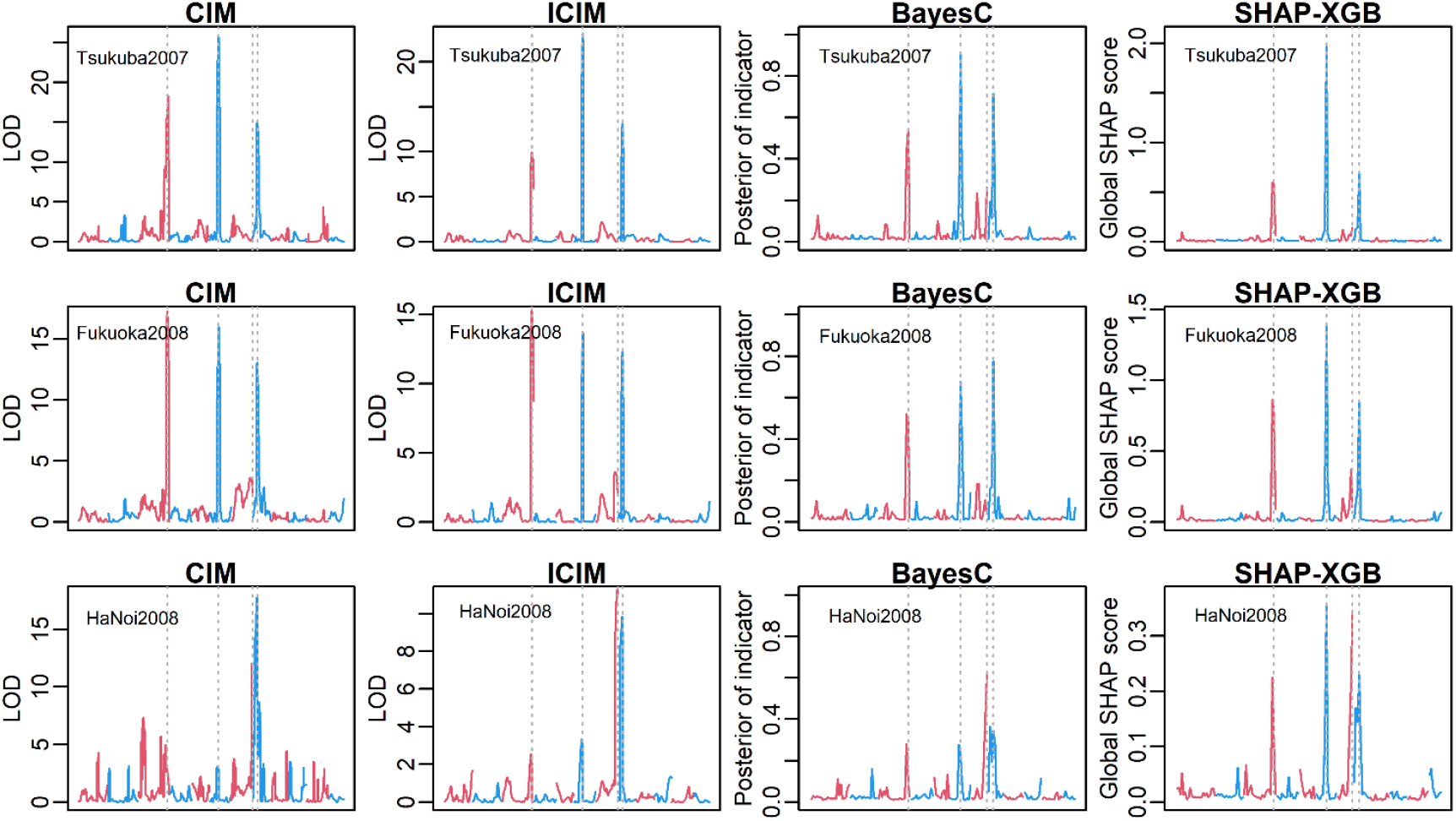
Comparison of QTL mapping methods to detect main effects in the rice heading date data. The y-axis denotes the QTL signals (CIM and ICIM, LOD score; BayesC, posterior indicator variable; SHAP-XGB, global SHAP score). The x-axis denotes the chromosome positions. The neighboring chromosomes are distinguished using different colors. The vertical bars indicate the four major QTLs (left to right, *Hd6, Hd1, Hd2*, and *DTH8*). Note that, the x axes differed among methods because CIM and ICIM conducted interval mapping whereas BayesC and SHAP-XGB used only marker genotypes as covariates. Three representative environments (Tsukuba2007, Fukuoka2008, and HaNoi2008) are presented. The full results are shown in Figure S5. CIM: composite interval mapping, ICIM: inclusive CIM, SHAP-XGB: Shapley additive explanations-assisted XGBoost.

**Fig. 6.**
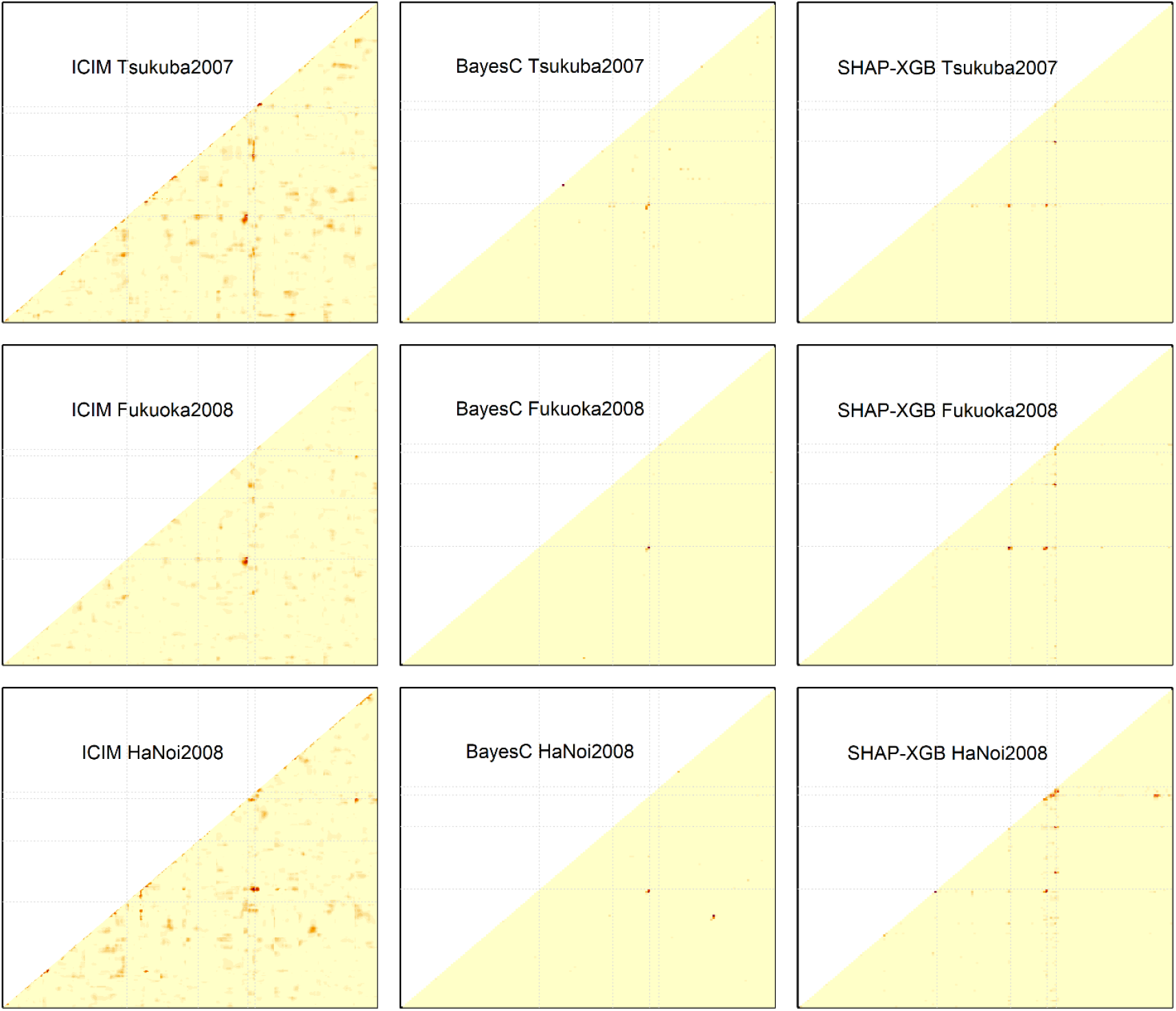
Comparison of QTL mapping methods to detect interaction effects in the rice heading date data. The y- and x-axes denote the relative positions of markers. The signal levels obtained with each QTL mapping method (ICIM: LOD score; BayesC: posterior indicator variable; SHAP-XGB: global SHAP interaction score) are shown using warm colors. The gray broken lines indicate the four major QTLs (left to right or bottom to top, *Hd6, Hd1, Hd2*, and *DTH8*). Note that positions of major QTLs in the x-axis slightly varied among methods because CIM and ICIM used interval mapping, whereas BayesC and SHAP-XGB used only marker genotypes as covariates. Three representative environments (Tsukuba2007, Fukuoka2008, and HaNoi2008) are presented. The full results are shown in Figure S6. ICIM: inclusive composite interval mapping, SHAP-XGB: Shapley additive explanations-assisted XGBoost.

**Fig. 7.**
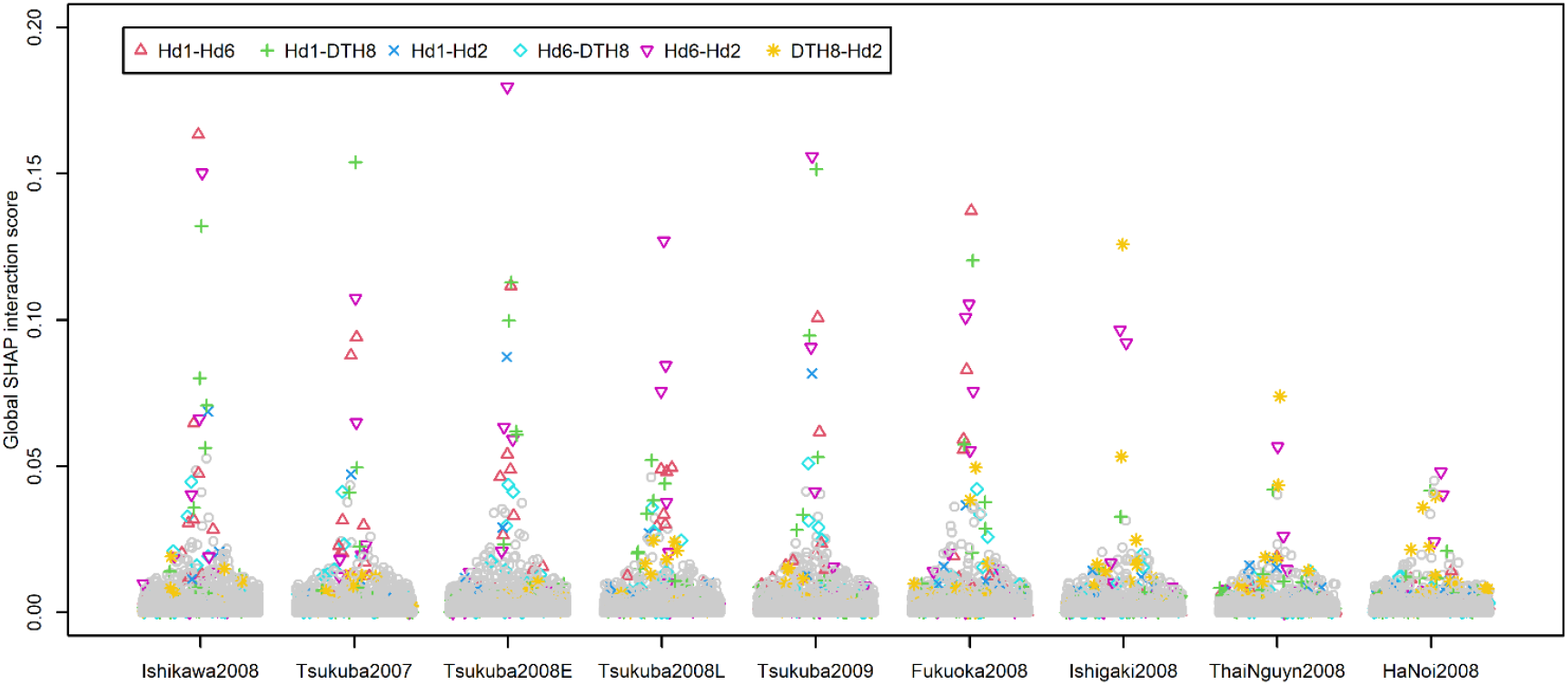
Summary of the global Shapley additive explanations (SHAP) interaction scores obtained in the rice heading date data. The y-axis denotes the global SHAP interaction scores, which are the averages of the absolute SHAP interaction values across all lines. The global SHAP interaction scores were horizontally jittered according to the densities of scores. The interaction values for the combinations of markers that were located within 20 cM from the known QTLs (*Hd6, Hd1, Hd2*, and *DTH8*) were regarded as the signals for the corresponding QTL interactions and highlighted using different colors and point shapes. The interactions between neighboring markers (< 20 cM apart) were removed for ease of visualization.

The interaction between *Hd1* and three QTLs (*DTH8, Hd6*, and *Hd2*) in Tsukuba2009 and the interaction between *DTH8* and *Hd2* in Ishigaki2008 were visualized using dependence plots (Fig. 8). The Koshihikari allele (1) of *DTH8* and *Hd6* tended to shorten days to heading at Tsukuba2009 when the Koshihikari alleles were harbored at *Hd1*, whereas the Kasalath allele (−1) tended to prolong days to heading when the Kasalath alleles were harbored at *Hd1*. Contrasting tendencies were observed for the interactions between *Hd1* and *Hd2* in Tsukuba2009 and *Hd2* and *DTH8* in Ishigaki2008.

**Fig. 8.**
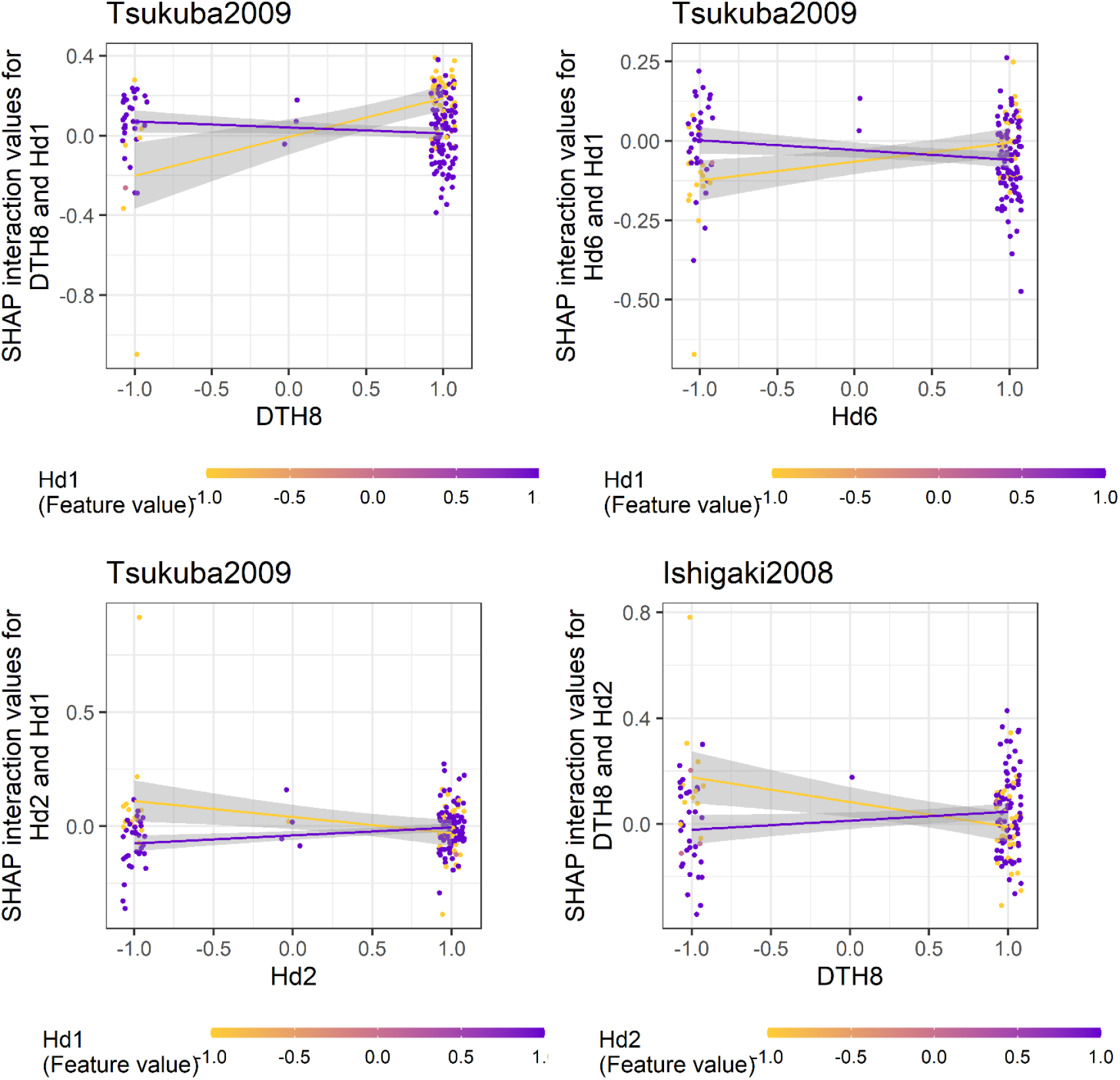
Dependencies between major QTLs shown with the Shapley additive explanations (SHAP) interaction values. The left top, right top, and left bottom panels show the dependency between *Hd1* and the three QTLs (*DTH8, Hd6*, and *Hd2*) at Tsukuba2009, respectively. The right bottom panel shows the dependency between *DTH8* and *Hd2* at Ishigaki2008. Each dot represents the SHAP interaction values for each line. Marker genotypes were regarded as the QTL genotypes. These markers were selected when the corresponding marker combinations showed the highest global SHAP interaction scores for the QTL interactions. For each panel and each QTL (marker), 1 and −1 indicate the Koshihikari and Kasalath alleles, respectively. The gray areas indicate the 95% confidence intervals of regression lines.

## Discussion

We applied SHAP-assisted XGBoost (SHAP-XGB) to QTL mapping in biparental populations and compared it with conventional mapping methods in this study. SHAP-XGB generally showed comparable accuracy in finding QTL main effects with conventional methods and higher accuracy in finding epistatic QTL effects. The dependency plots enabled easy interpretation of the interactions. These results suggested that SHAP-XGB complements existing methods, particularly for identifying novel interactions between QTLs.

The simulation results revealed several properties of the SHAP-XGB. First, compound and pure global SHAP scores worked similarly to map the main effects of QTLs. While the compound scores showed slightly increased detection power when epistasis variances were larger, they also showed a slight increase in false positives when QTLs with interaction effects lacked the main effects. Thus, using compound scores may be advantageous as long as QTLs with non-zero main effects are involved in the interactions. Using compound scores may circumvent the reduction in pure scores due to LD, as described later. Second, the degree of fit (i.e., the extent to which the model fits the data) of the XGBoost models did not affect the QTL detection power. This is because the ROC curves and AUC values were calculated based on the relative rankings of markers that did not differ significantly depending on the fitting status. However, the result of over- or underfitting is the inflation or deflation of SHAP values, which in turn may make the estimation of QTL effect sizes difficult. Thus, careful tuning of the fitting degree is required to improve interpretability. Finally, SHAP-XGB is robust to the misspecification of target effects, i.e., additive or dominance. That is, even when SHAP-XGB targets additive effects, it can map dominance effects or dominance-by-dominance interaction effects as accurately as the existing methods with correct target specifications. This property may be advantageous; however, it may be disadvantageous in some cases because the gene actions of the detected QTLs cannot be identified. To capture the additive and dominance genetic effects separately, it is possible to incorporate both the additive and dominance genetic coding of a single marker concurrently in SHAP-XGB. However, such modeling would not work as expected because the additive coding in SHAP-XGB would capture the dominance effects and the QTL signals would be diluted. This can be illustrated as follows. Suppose there is a QTL influencing a trait, and its gene action is overdominance, resulting in phenotypic values of 0 (homozygote), 1 (heterozygote), and 0 (the other homozygote), ignoring environmental effects. If XGBoost is applied to these data using SNPs encoded additively (0, 1, and 2) as features, it will attempt to reduce the residuals by stacking decision trees. Assume the first tree is constructed with a rule where genotype 0 of SNP *A* (a SNP linked to the QTL) is assigned a value of 0, and genotypes 1 and 2 are assigned values of 1. The resulting residuals become 0 (homozygote), 0 (heterozygote), and −1 (homozygote). Next, if a second tree is built to further reduce the residuals using a rule where genotypes 0 and 1 of SNP A are assigned 0, and genotype 2 is assigned −1, the model achieves a perfect fit to the data.

There are ways to measure feature importance for gradient boosting other than SHAP, including “gain” (the average increase of the objective function when the feature is added to trees) and “split count” (the number of times that trees split using the feature). Although these indices are provided by the R package of XGBoost, we did not adopt them because they do not hold the consistency property (Lundberg et al. 2018) and the package did not support them as importance of feature interactions. Nevertheless, considering that there are various indices to measure feature importance, such as local interpretable model-agnostic explanations (LIME) (Ribeiro et al. 2016) and the Shapley–Taylor interaction index (Sundararajan et al. 2020), it would still be worth investigating which indices are suitable for QTL mapping and whether the properties revealed here are applicable when SHAP is replaced with other indices. Such investigations are required to mitigate the negative effects of LD, as will be discussed later.

Although the simulated and real data analyses consistently suggest the usefulness of SHAP-XGB in QTL mapping, the marker genotypes have an undesirable property, i.e., LD between markers, for evaluating feature importance. Because XGBoost randomly samples markers to build a tree *f*, QTL effects are diluted with the markers linked to the QTLs, consequently decreasing the Shapley values of each linking marker. In fact, the global SHAP scores and SHAP interaction values in Figs. 5 and 7 shrank to 0 (2.0 at most) considering that the units of the scores/values are the same as the phenotypic values (i.e., days). These decreased values were probably due to the correlated marker genotypes. The LD between the markers affected another aspect. In rice heading date analyses, interactions between neighboring markers were occasionally assigned large SHAP interaction values, resulting in a reduction in the main pure scores (these interactions were removed from Fig. 7 for ease of visualization). However, such interactions are caused by LD between neighboring markers and thus should be interpreted as part of the main QTL effects. Although this decrease in pure main effects can be circumvented using compound scores for mapping, caution is required when interpreting pure scores. It has also been pointed out that approximations of Shapley values with SHAP become inaccurate when features are correlated (Aas et al. 2021) because the evaluations of the expectation in Eq. (2) assume independence between the features. Thus, taken together, LD would affect the detection power of SHAP-XGB. One solution is to replace SHAP with other indices that consider the correlations among features (Aas et al. 2021).

Interactions between the major heading date QTLs weakened at lower latitudes (Fig. 7 and Table 2). These results were reasonable because *Hd6, Hd2*, and *DTH8* are involved in heading date variation under long-day conditions (Hori et al. 2016, Wei et al. 2020). Interactions between *Hd1* and three QTLs (*Hd6, Hd2*, and *DTH8*) are plausible under natural long-day conditions. For example, interactions between functional *Hd6* alleles (e.g., Kasalath allele) and *Hd1* alleles (e.g., Nipponbare allele) were reported under long-day conditions (Ogiso et al. 2010). Interactions between *Hd1* and *Hd2* under long-day conditions (Lin et al. 2000), and interactions among *Hd1, Hd2, DTH8*, and *Ghd7* under natural long-day conditions were also reported (Zhang et al. 2019). The interactions between *Hd2* and *DTH8*, which is prominent in lower latitudes, are interesting but also plausible because interactions between the functional/non-functional alleles of *Hd2* and *DTH8* under natural short-day conditions have been reported (Zhang et al. 2019). Thus, these results indicated the usefulness of epistatic QTL mapping using SHAP-XGB for identifying novel interactions.

SHAP-XGB may be applicable to studies with denser DNA markers, e.g., GWAS; however, the computational time and required memory size will increase. The increase in computational time depends on the XGBoost fitting and SHAP value calculations. Fitting XGBoost to the rice real data (*n* = 176 and *p* = 162) took 0.111 ± 0.055 seconds using a machine with Intel Corei7-13700k (3.40 GHz), 24 logical cores, and 128GB RAM, and calculation of SHAP and SHAP interaction scores took 0.007 ± 0.014 and 0.316 ± 0.144 seconds, respectively. We generated a feature matrix with *n* = 1,760 and *p* = 162,000 from the standard normal distribution and fitted the SHAP-XGB to investigate the scalability of SHAP-XGB. XGBoost took 51.34 seconds, and the calculation of SHAP took 430.89 seconds using the same machine, which is sufficiently practical. However, the calculation of the SHAP interaction scores failed because of a memory allocation error, and this error could not be fixed until *p* decreased to 2,000. SHAP interaction scores are calculated for all marker combinations in all samples (i.e., an *n* × *p* × *p* array is created), and the memory requirement may be a major bottleneck. Thus, it may be necessary to prune markers using, for example, t-tests or sure independence screening (Fan and Lv 2008). In GWAS, population stratification will be another issue. One solution would be to adjust the phenotypic values for population stratification before application.

Because SHAP-XGB outperformed linear statistical methods in mapping QTL interactions, it is likely that SHAP-XGB and other machine learning methods are more suitable than linear methods for mapping higher-order and/or complex interactions among QTLs. Although the present study addressed only two-way interactions, this issue is worth investigating. Another possible extension of SHAP-XGB is QTL mapping for binary or categorical traits, because XGBoost can be applied to classification problems.

The lack of thresholds for determining significant markers is a practical drawback for each application of SHAP-XGB (QTL mapping and GWAS). Permutation tests are one possible solution. Permutation tests are generally used to determine the thresholds for IM and CIM (e.g., Broman et al. 2003) and have been applied to detect gene-gene interactions based on the Shapley-based interaction score and neural networks (Cui et al. 2022). However, further investigation is required to determine whether these tests work as expected.

## Author contribution statement

TI and AO collaborated in analyzing the data and drafting the manuscript. AO conceived the study.

## Acknowledgements

The authors would like to thank Ryukoku University for providing financial support.

