## Supporting Information for "Applying gradient tree boosting to QTL mapping with Shapley additive explanations"

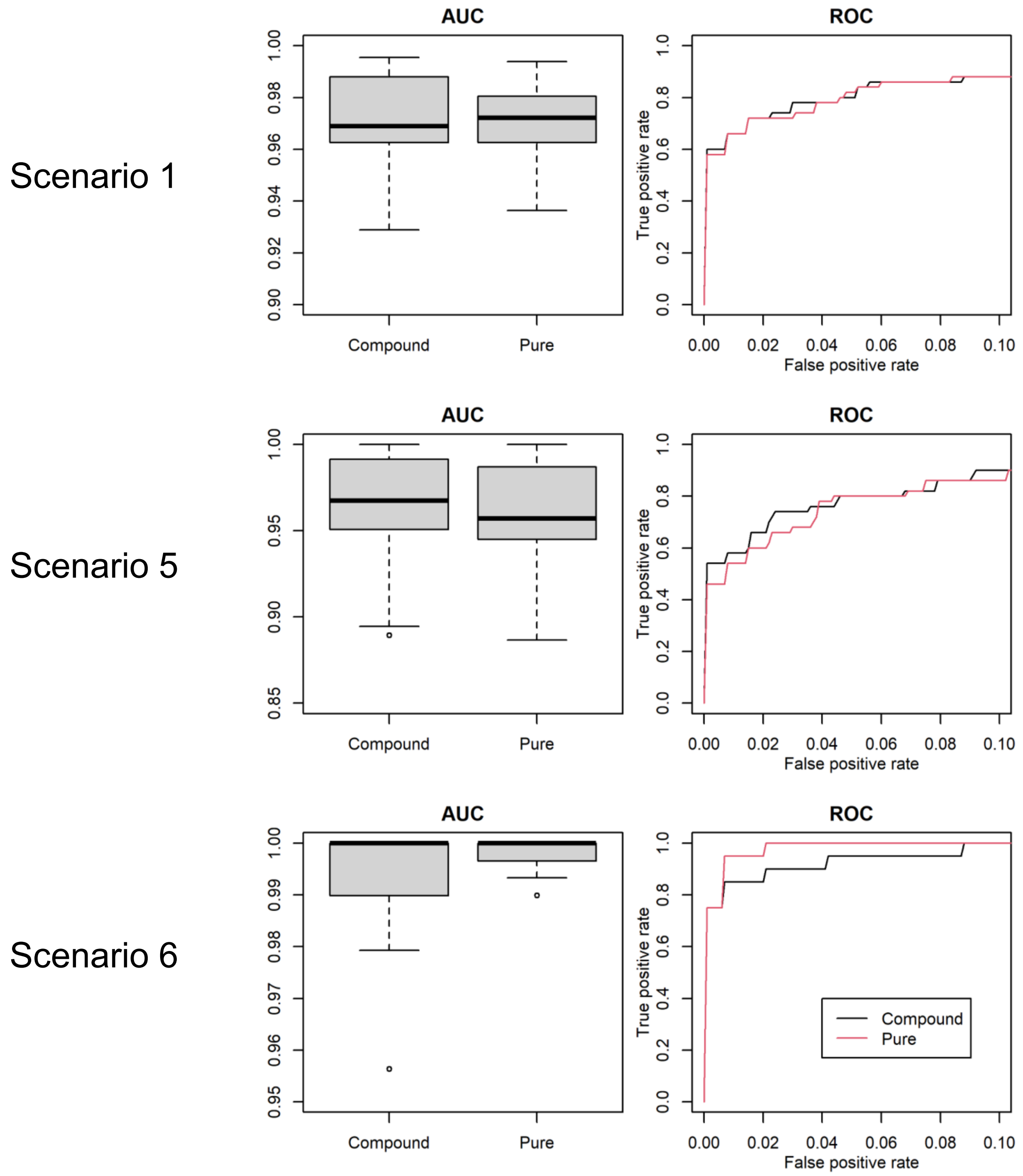


**Supplemental Fig. 1** Comparison of area under the curve (AUC) values and receiver operating characteristic (ROC) curves between compound and pure global SHAP scores. AUC values and ROC curves were evaluated in the three simulation scenarios, 1, 5, and 6. The ROC curves are the mean curves of 10 replications.

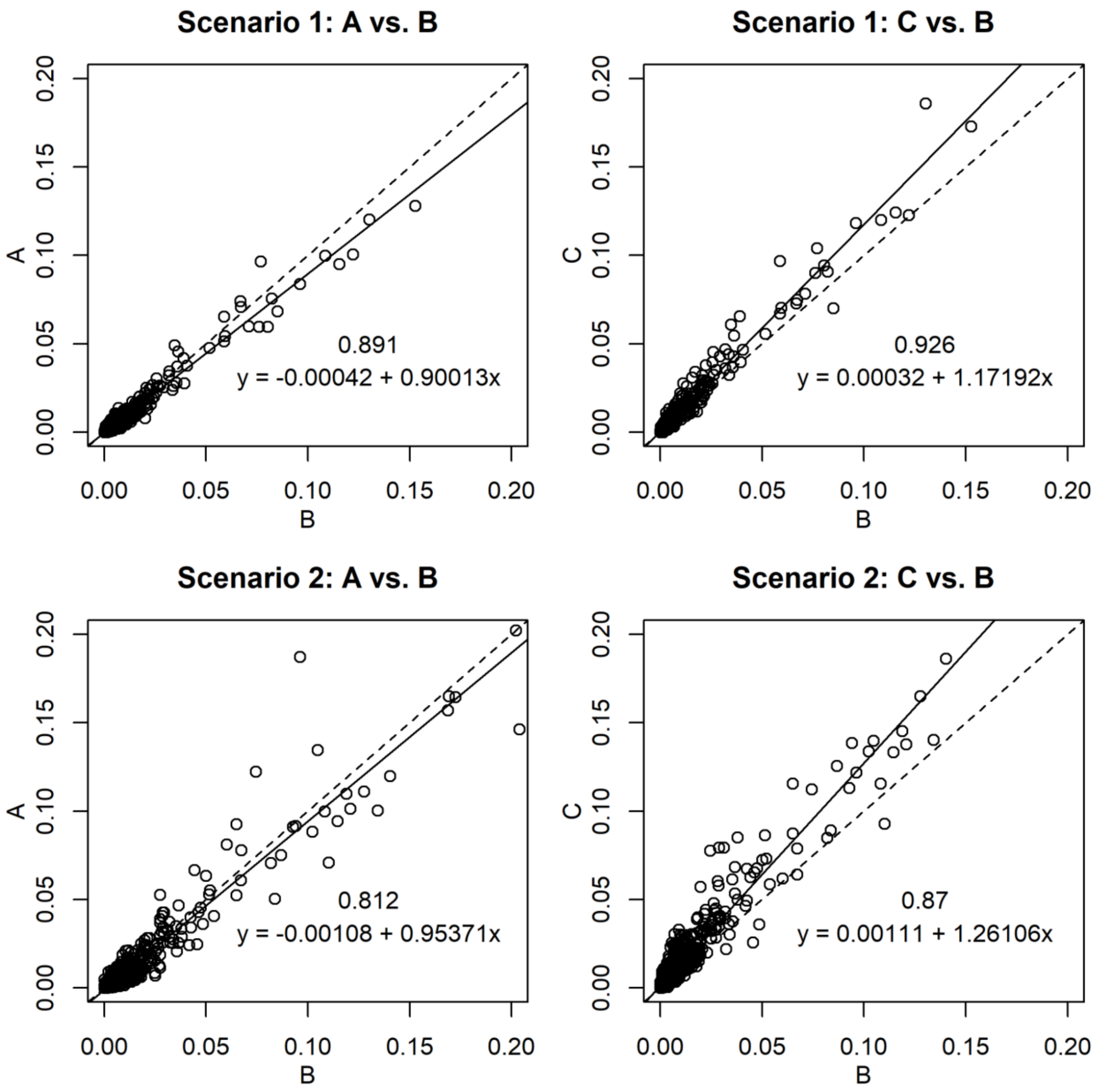


**Supplemental Fig. 2** Comparison of the compound global SHAP scores obtained using the different XGBoost settings. A, B, and C denote the XGBoost settings resulting in underfitting, moderate, and overfitting, respectively. The global SHAP scores were the averages of absolute SHAP values across all lines. The regression lines are shown with solid lines and the 1:1 lines are with broken lines. Numbers in the plots are Spearman correlation coefficients and equations are regression equations.

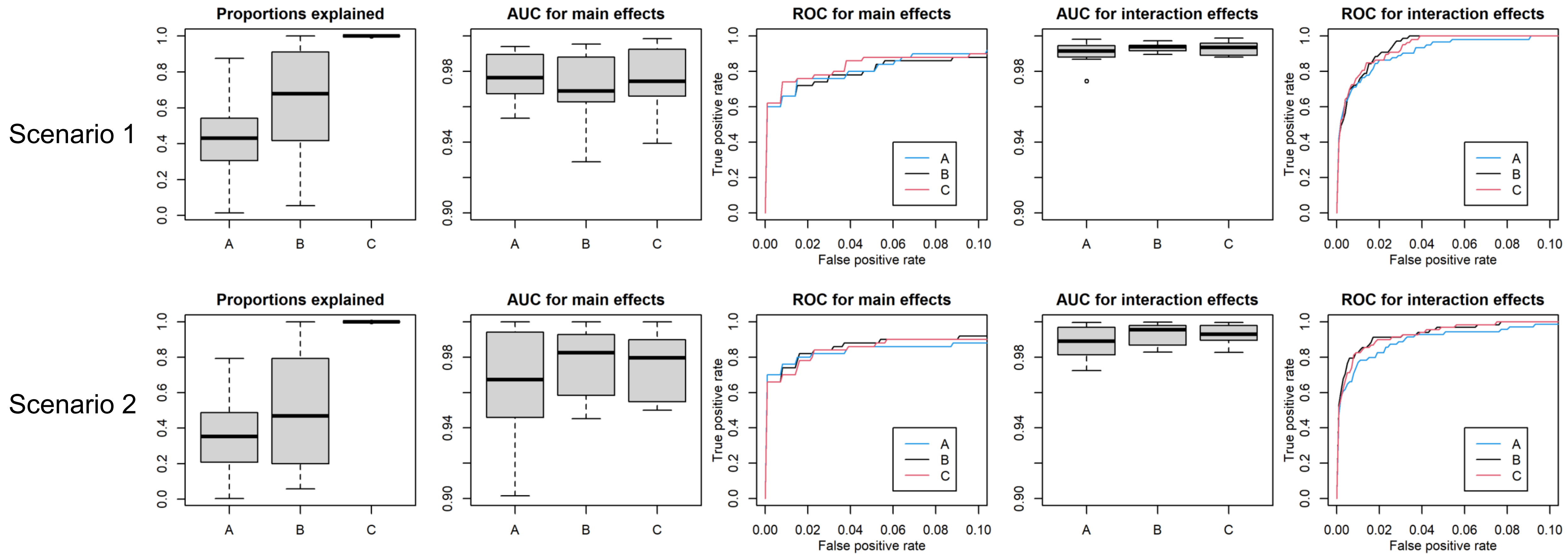


**Supplemental Fig. 3** Comparison of XGBoost settings in the simulated data sets (scenarios 1 and 2, top and bottom, respectively). A, B, and C denote the XGBoost settings resulted in underfitting, moderate fitting, and overfitting, respectively. The left panels show proportions of phenotypic variances explained by the models. The other panels show the area under the curve (AUC) values and the receiver operating characteristic (ROC) curves for the main and interaction effects of QTLs. ROC curves are the mean curves of 10 replications.

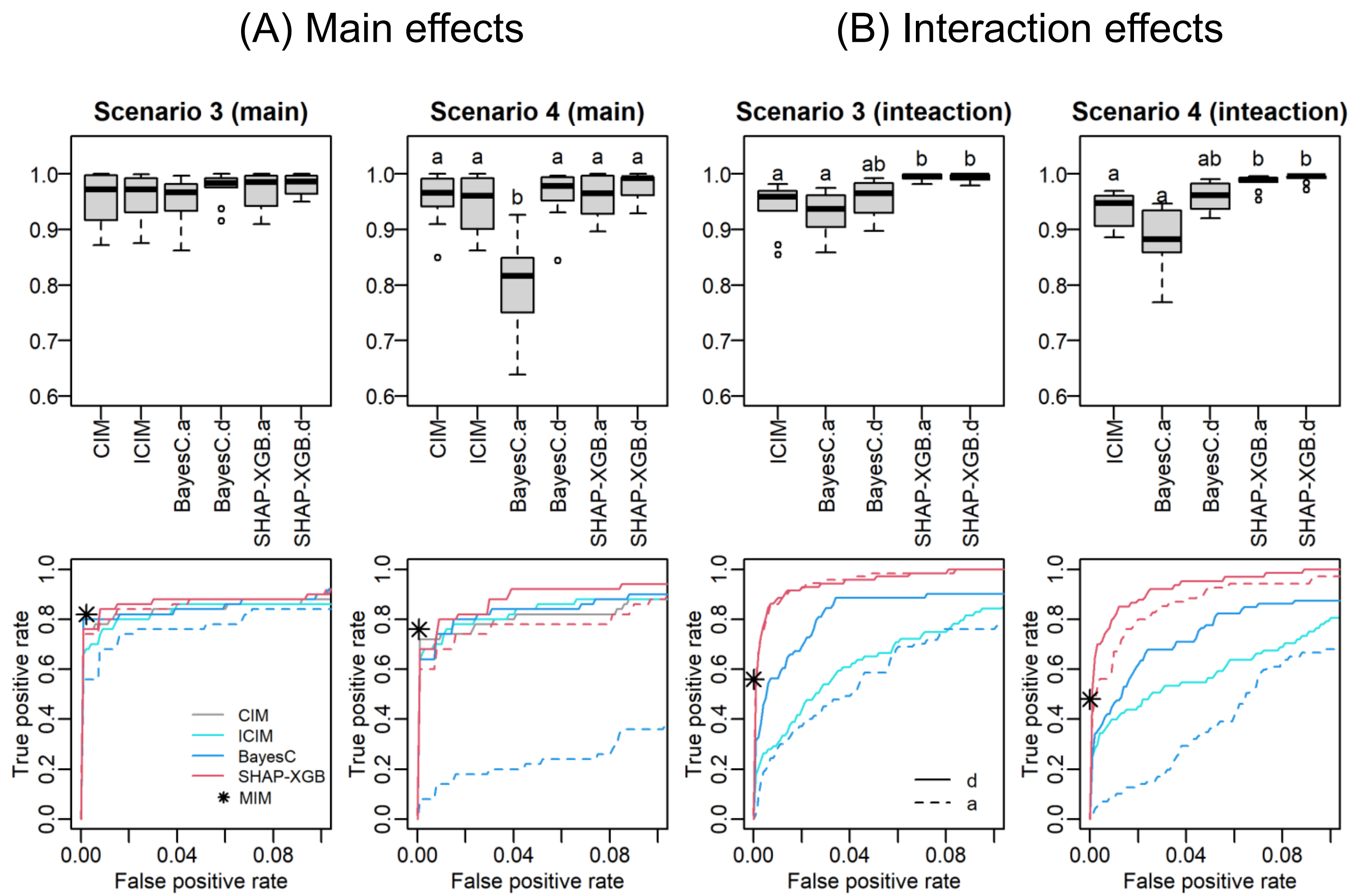


**Supplemental Fig. 4** Area under the curve (AUC) values and receiver operation characteristic (ROC) curves in mapping main (A) and interaction (B) effects when dominance effects were simulated. The top and bottom panels are the AUC values and ROC curves, respectively. The ROC curves are the mean curves of 10 replications. Scenarios 3 and 4 simulated $F_2$ populations that comprised only complete and overdominance effects, respectively. Small letters a and d indicate methods that targeted the additive and dominance effects, respectively. Different letters in the box plots denote significant differences ($P$ <0.05). Because the results of MIM are the lists of detected QTLs and their interactions, true and false positives are shown with the asterisk in the bottom plots. CIM: composite interval mapping, ICIM: inclusive CIM, SHAP-XGB: Shapley additive explanations-assisted XGBoost, MIM: multiple interval mapping.

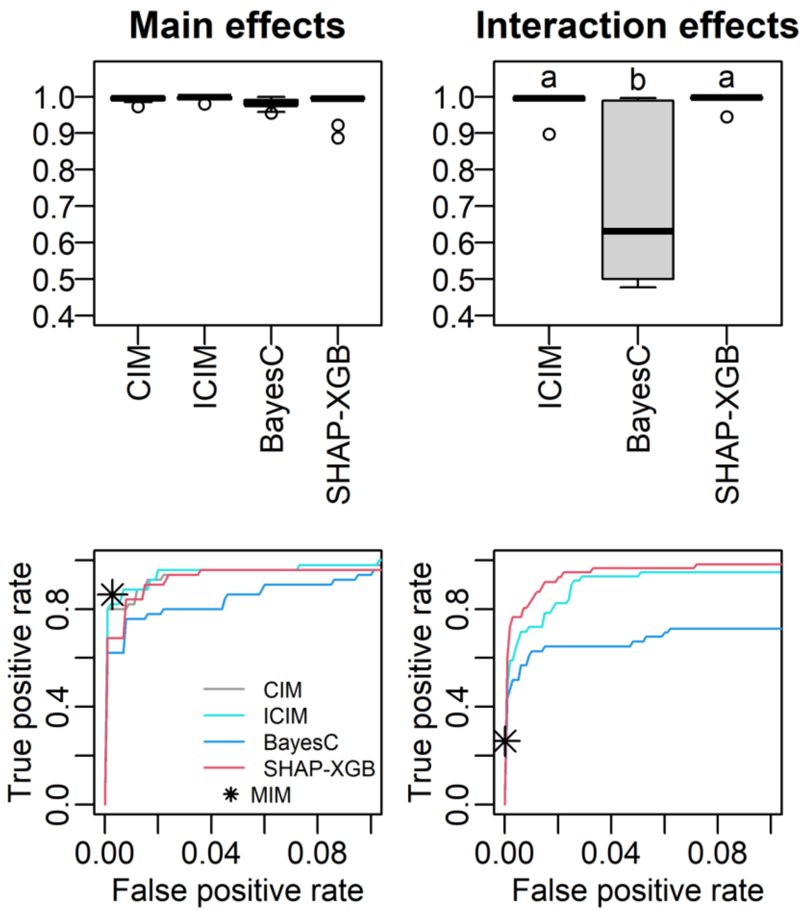


**Supplemental Fig. 5** Area under the curve (AUC) values and receiver operation characteristic (ROC) curves in mapping main and interaction effects when QTLs were linked to each other (Scenario 9). The top and bottom panels are the AUC values and ROC curves, respectively. The ROC curves are the mean curves of 10 replications. Different letters in the box plots denote significant differences (*P* <0.05). Because the results of MIM are the lists of detected QTLs and their interactions, true and false positives are shown with the asterisk in the bottom plots. CIM: composite interval mapping, ICIM: inclusive CIM, SHAP-XGB: Shapley additive explanations-assisted XGBoost, MIM: multiple interval mapping.

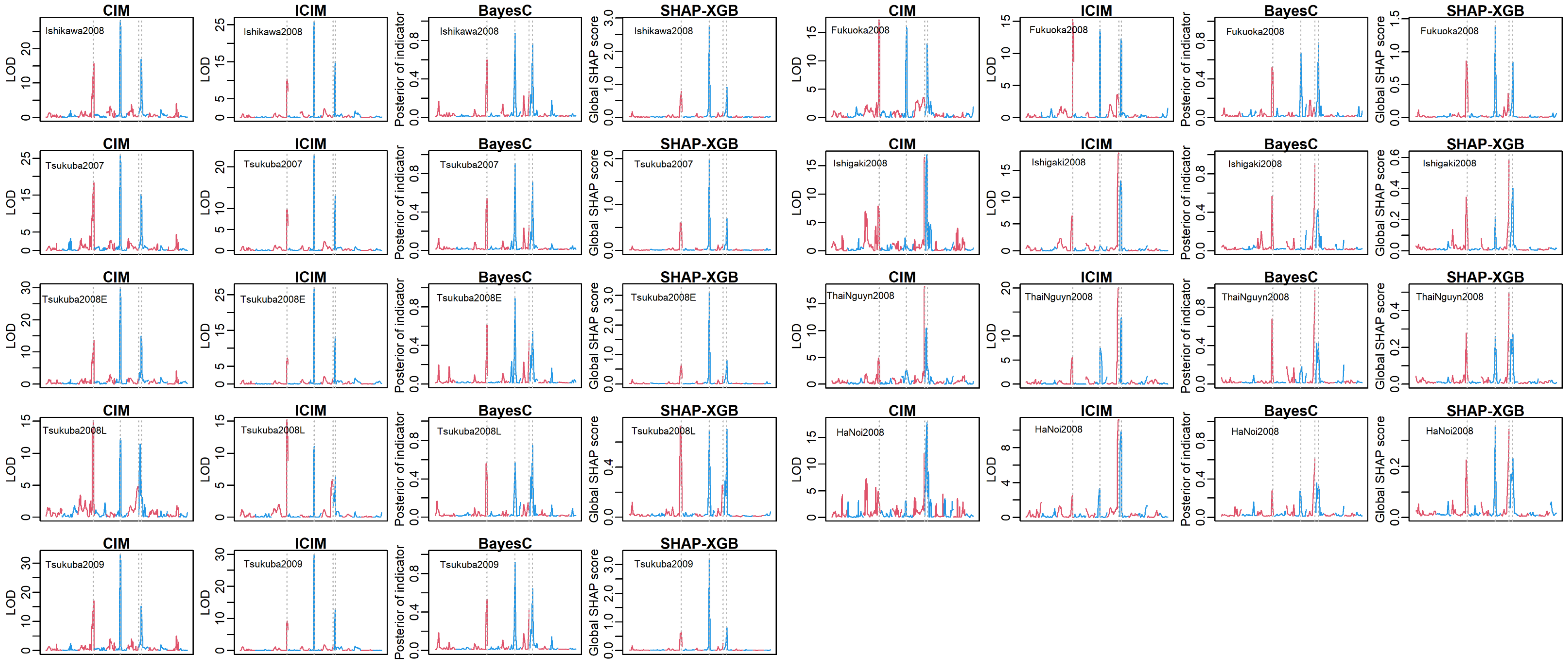


**Supplemental Fig. 6** Comparison of QTL mapping methods to detect main effects in the rice heading date data. The y axis denotes the QTL signals (CIM and ICIM, LOD score; BayesC, posterior of indicator variable; SHAP-XGB, global SHAP score). The x axis denotes the chromosome positions. The neighboring chromosomes are distinguished with different colors. The vertical bars indicate the four major QTLs (left to right, *Hd6*, *Hd1*, *Hd2*, and *DTH8*, respectively). Note that, because CIM and ICIM conducted interval mapping whereas BayesC and SHAP-XGB used only marker genotypes as covariates, x axes differed among methods. CIM: composite interval mapping, ICIM: inclusive CIM, SHAP-XGB: Shapley additive explanations-assisted XGBoost.

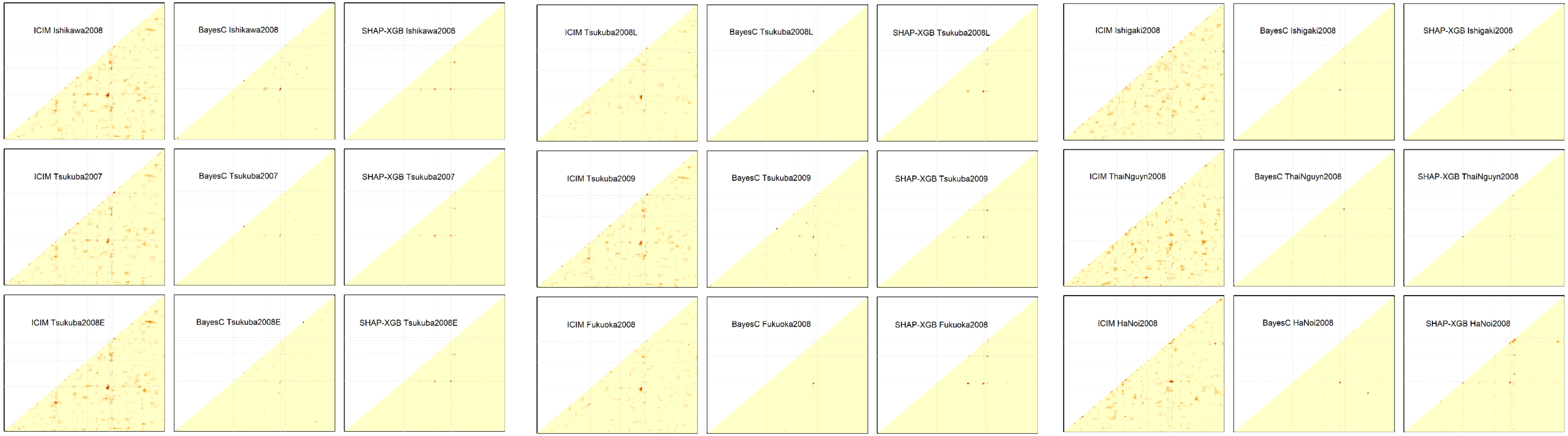


**Supplemental Fig. 7** Comparison of QTL mapping methods to detect interaction effects in the rice heading date data. The y and x axes denote the relative positions of markers. The signal levels obtained with each QTL mapping methods (ICIM, LOC score; BayesC, posterior indicator variable; SHAP-XGB, global SHAP interaction score) are shown with heat colors. The gray broken lines indicate the four major QTLs (left to right or bottom to top, *Hd6*, *Hd1*, *Hd2*, and *DTH8*, respectively). Note that, because CIM and ICIM conducted interval mapping whereas BayesC and SHAP-XGB used only marker genotypes as covariates, positions of major QTLs in the x axes slightly differed among methods. ICIM: inclusive composite interval mapping, SHAP-XGB: Shapley additive explanations-assisted XGBoost.

**Supplemental Table 1** QTLs and QTL interactions detected with multiple interval mapping in the real rice data

| Environment | Main effects | | | | Interact with | | | |
|---|---|---|---|---|---|---|---|---|
| | Chr | Position (cM) | Major gene | Size | Chr | Position (cM) | Major gene | Size |
| Ishikawa2008 | 3 | 95.33 | *Hd6* | 0.6362 | | | | |
| | 5 | 68.73 | | 2.2831 | | | | |
| | 6 | 30.57 | *Hd1* | -7.9966 | | | | |
| | 7 | 91.5 | *Hd2* | -1.6585 | | | | |
| | 8 | 18.73 | *DTH8* | 8.0201 | | | | |
| | 11 | 3.85 | | 1.1193 | | | | |
| Tsukuba2007 | 1 | 35.44 | | 1.4103 | | | | |
| | 3 | 94.45 | *Hd6* | 0.6988 | | | | |
| | 5 | 68.73 | | 1.8493 | | | | |
| | 6 | 30.57 | *Hd1* | -6.0632 | | | | |
| | 7 | 91.5 | *Hd2* | -1.1049 | | | | |
| | 8 | 22.54 | *DTH8* | 5.7596 | | | | |
| Tsukuba2008E | 3 | 96.19 | *Hd6* | 0.1404 | | | | |
| | 5 | 68.73 | | 1.8944 | | | | |
| | 6 | 30.57 | *Hd1* | -8.7395 | | | | |
| | 7 | 91.5 | *Hd2* | -2.6196 | | | | |
| | 8 | 22.54 | *DTH8* | 7.8378 | | | | |
| | 11 | 3.85 | | 0.6623 | | | | |
| Tsukuba2008L | 3 | 122.63 | *Hd6* | 3.6521 | 7 | 91.5 | *Hd2* | -2.753 |
| | 6 | 30.57 | *Hd1* | -3.4591 | | | | |
| | 7 | 25.8 | | 1.3684 | | | | |
| | 7 | 91.5 | *Hd2* | 0.8667 | | | | |
| | 8 | 17.83 | *DTH8* | 5.2241 | | | | |
| | 11 | 3.85 | | 0.1474 | | | | |
| Tsukuba2009 | 3 | 98.67 | *Hd6* | 0.5534 | | | | |
| | 5 | 68.73 | | 2.1994 | | | | |
| | 6 | 30.57 | *Hd1* | -8.18 | | | | |
| | 7 | 91.5 | *Hd2* | -2.1135 | | | | |
| | 8 | 18.73 | *DTH8* | 7.4722 | | | | |
| | 11 | 1.97 | | 1.0456 | | | | |

(Table S1 continued)

| Environment | Main effects | | | | Interact with | | | |
|---|---|---|---|---|---|---|---|---|
| | Chr | Position (cM) | Major gene | Size | Chr | Position (cM) | Major gene | Size |
| Fukuoka2008 | 1 | 35.43 | | 1.2397 | | | | |
| | 3 | 124.69 | *Hd6* | 2.9079 | 6 | 30.57 | *Hd1* | -3.026 |
| | | | | | 7 | 83.59 | *Hd2* | -2.977 |
| | 6 | 30.57 | *Hd1* | -6.5639 | | | | |
| | 7 | 25.89 | | 1.616 | | | | |
| | 7 | 83.59 | *Hd2* | 1.584 | 8 | 18.72 | *DTH8* | 2.435 |
| | 8 | 18.72 | *DTH8* | 6.3669 | | | | |
| Ishigaki2008 | 1 | 35.44 | | -5.3063 | 11 | 3.85 | | -6.482 |
| | 3 | 117.23 | *Hd6* | 2.1597 | 7 | 91.5 | *Hd2* | -1.565 |
| | 6 | 13.02 | | -0.3499 | | | | |
| | 7 | 91.5 | *Hd2* | 5.0085 | 8 | 13.07 | *DTH8* | 2.529 |
| | 8 | 13.07 | *DTH8* | 4.6862 | | | | |
| | 11 | 3.85 | | -5.8289 | | | | |
| ThaiNguyn2008 | 1 | 4.77 | | -2.4888 | | | | |
| | 3 | 117.23 | *Hd6* | 1.8787 | | | | |
| | 6 | 12.03 | | 0.743 | | | | |
| | 7 | 91.5 | *Hd2* | 4.4247 | | | | |
| | 8 | 15.03 | *DTH8* | 3.4037 | | | | |
| | 11 | 0.01 | | -1.1966 | | | | |
| HaNoi2008 | 3 | 96.19 | *Hd6* | 4.9436 | 8 | 22.54 | *DTH8* | 5.721 |
| | 6 | 30.57 | *Hd1* | 2.474 | | | | |
| | 7 | 86.39 | *Hd2* | 2.4728 | | | | |
| | 8 | 22.54 | *DTH8* | 2.2514 | 10 | 46.71 | | -4.793 |
| | 10 | 46.71 | | -4.8473 | | | | |
| | 11 | 3.85 | | 0.3141 | | | | |
