## Supplementary Text for "Applying gradient tree boosting to QTL mapping with Shapley additive explanations"

### Reviewers’ comments for versions 1-5

### -The current (final) version is 7

### Decision letter (for ver. 1)

### Theoretical and Applied Genetics

Dear Prof. Onogi,

Thank you for submitting your manuscript to TAG. Your manuscript TAAG-D-24-00051 "Applying gradient tree boosting to QTL mapping with Shapley additive explanations" was evaluated by external reviewers, and their comments are given below. You are kindly requested to also check the website for possible reviewer attachment(s).

I am sorry to inform you that your manuscript cannot be accepted for publication in TAG in the present form. Although TAG is rejecting the present manuscript, a thoroughly revised manuscript addressing all the reviewers' suggestions could be suitable for publication in TAG. However, a revised manuscript will be handled as a new submission that can again enter the review process. A new submission must refer to the present manuscript number and you will need to highlight the changes you made in the manuscript as well as provide detailed and itemized responses to the comments of the reviewers and Editor.

On behalf of the journal, we thank you for submitting your manuscript to TAG and hope that the outcome of this specific review will not discourage you from sending future papers to TAG.

With kind regards,
Antonio Augusto Franco Garcia
Editor
Theoretical and Applied Genetics

#### Editor:

Your comparison with other methods is very biased and unfair. Nobody should use any of the mentioned software using the defaults parameters, but considering the ongoing data set and cenario.

This must be considered and fixed if you want your paper to have another round of evaluations.

Moreover, MIM is your major reference point, since it is the most relevant method for QTL mapping including epistasis. CIM is well known for not allowing apropriate estimation of epistasis, as discussed in the papers proposing MIM. Also, currently MIM should be done in R/qtl (with apropriate parameters), not on QTLCartographer, which is very outdated. BayesC, etc, are not designed for mapping QTL, but for doing genomic selection, so using it in a wrong cenario is also unfair.

Without these points I will not consider a ressubmission.

#### Reviewer #1:

This study evaluates the potential of a gradient boosting algorithm based called SHAP-XGB to identify epistasis in simulated and real (rice) biparental mapping populations. SHAP-XGB was compared to existing approaches, and it was shown that SHAP-XGB identified more epistatic loci than conventional methods, such as CIM, MIM, and BayesC, while having equivalent detection rates for main-effect loci.

concepts from a simple, intuitive perspective (the analogy of using Shapley values with contributions from team players was very helpful); as well as providing technical detail. The Results were presented very clearly, and the Discussion touched upon some really important points.

Major Points

1.) The ability to detect epistasis plays a prominent role in the work presented in this paper. I suggest editing the Abstract so that the word "epistasis" is used at least once.

2.) Question out of curiosity: To quote Lines 310-311, "SHAP was originally proposed as a local importance measure". When translating this concept to quantitative genetics, does this mean that SHAP is geared (I don't that "bias" the correct word to use here) towards detecting epistatic interactions between loci that are in LD and/or physical proximity to each other?

3.) Misspecification - I think it would be informative to report from the simulations studies how often SHAP misspecifies, e.g., additive QTNs as epistatic, etc. This is touched upon in the Results, but would it be possible to dedicate a specific section of the Results to this?

Minor Points

Lines 81-86: This is an excellent introduction to SHAP. I really like how it steers away from technical jargon. However, there is important things missing - I couldn't figure out what "instance" is referring to. Could some text be added here to briefly introduce the reader to what is meant by "instance"? (I saw that the abstract defines an instance of a "plant/line", but this should be stated again)

Line 92 - please describe what XGBoost here.

Line 203 - please change "LTR" to "LRT"

Line 274 - could some clarification please be provided for compiling the MIM results (where an ROC curve was not used). For example, was a permutation procedure used to obtain genome-wide significance thresholds at ¥alpha = 0.05?

Line 332 - I suggest not referring these to e.g., "Scenarios 1, 5, and 6" but instead as what is being simulated (e.g. F_2 lines with additive effects and additive-by-additive epistasis). The reader will then not need to stop reading and refer back to Table 1

Figure 7: What do the grey areas around the purple and yellow lines depict? Prediction intervals? In any case, please indicate what these are in the caption.

Line 460 - "We applied SHAP-XGB and SHAP-assisted XGBoost": this is a typo, right? If I understand correctly, SHAP-XGB is the acronym for SHAP-assisted XGBoost.

Line 472 - what is meant by "degree of fit"?

Line 478-480 - Re: misspecification. Is this saying that, e..g., SHAP-XGB can identify dominance main effects with a high true positive rate but they are misspecified as additive? If yes, maybe rephrase this to convey this concept.

#### Reviewer #2:

I have read the present manuscript submitted to Theoretical and Applied Genetics (TAAG). The study aimed to apply a machine learning-based extreme gradient tree boosting method (XGBoost) for quantitative trait loci (QTL) mapping and compare it with traditional approaches, such as likelihood ratio test (LRT), composite interval mapping (CIM), multiple interval mapping (MIM), and BayesC. For this purpose, authors have used: (i) simulated data from two experimental populations – an F2 and a recombinant inbred line (RIL) – considering six distinct scenarios regarding the effects of pre-defined QTL; and (ii) real data from a rice (Oryza sativa L.) backcross experimental population (BC) consisting of 174 individuals, which were previously genotyped with 162 restriction fragment length polymorphism (RFLP) markers and phenotyped for heading date in nine environments from six field trials.

In general, the authors were motivated to evaluate whether XGBoost, assisted by the Shapley additive explanations (SHAP) algorithm, could be helpful for mainly detecting complex interactions (epistasis) between QTL. In this context, they wondered whether SHAP-assisted XGBoost (SHAP-XGB) could complement traditional approaches and additionally contribute to QTL studies. In my opinion, the authors' motivation is valid and interesting. However, there are concerns regarding the strategies adopted by them, and I would like to present below relevant points related to these concerns. Authors are required to address each of these points, with the aim of improving their manuscript and make it acceptable for publication.

MAJOR POINTS:

1)

XGBoost is formally presented in the subsection "Extreme gradient boosting", within the section "Materials and Methods". How could the parameter/variable "q" affect the response of an instance (first equation), being the structure of a decision tree? Moreover, what would it be the parameter/variable "n" specified in the second equation? It is totally unclear for readers. Details from the XGBoost theory should be included in the text. Furthermore, I believe the authors could also include short sentences in the text containing the intuition behind all the equations presented, which would appropriately guide readers to understand the method. Certainly, these additional insights would significantly contribute to readers' understanding and enrich the manuscript;

2)

SHAP-XGB is presented in detail in the subsection "Algorithm of SHAP-XGB", also within the section "Materials and Methods". Although the authors have explained the method using a step-by-step procedure, I believe they could include an illustrative representation or scheme containing all these SHAP-XGB steps. This representation/scheme could be constructed and shaped considering the context of QTL mapping, following the authors' current purpose. In my opinion, the contribution would significantly be important to readers' understanding, in the same way as was suggested in the previous major point;

3)

In the subsection "Comparing methods", still within the section "Materials and Methods", authors mentioned that the SHAP-XGB method was compared with methods which were selected based on their use in QTL mapping studies and/or ease of execution in the R environment. What did the authors mean with the latter? Did they refer to the methods that are fully implemented and available in the R software via specific packages? Please, be careful with these statements. Readers could easily interpret that only methods with a user-friendly code routine have been selected to the present study. Of course, methods fully implemented in R could be very useful for the scientific community. However, their availability in the current software should not necessarily be related to their importance. Therefore, I would suggest the authors to remove this sentence and avoid using these arguments, so that users would not be encouraged to use methods via a simple "mouse click" even within R environment.

Still within the same subsection, it is mentioned that all methods only used the marker genotypes as covariates, so that loci genotype probabilities between markers (intervals) were not considered for QTL mapping. Why did the authors proceed like that? Have the authors performed QTL mapping without a previously built genetic linkage map for each of the simulated and real populations? Have they mapped QTL as a single marker analysis (SMA) even with the availability of the chromosome and the position of each marker? In this sense, how did they effectively perform both CIM and MIM, which rely on a previous genetic linkage map? I believe the authors have used a strategy to accurately compare the different methods selected. However, from my point of view, it is totally unclear for readers in the manuscript. Please, address this very relevant point from the study;

4)

Regarding the subsection "Likelihood ratio test (LRT)", I checked that the LRT method was proposed in the context of genome-wide association studies (GWAS). The method was fully implemented in the R package named SNPassoc. However, no details about the method was presented in the paper from this package. In this sense, where was the method formally published including the likelihood function or equation of the tested/evaluated models? Readers should be able to access all this information;

5)

Regarding the subsection “Composite interval mapping (CIM)”, why did the authors use a set of default parameters within the cim function from the qtl R package? Did these default parameters really make sense considering the context of this study? Sincerely, I wonder if the use of such default parameters was really associated with a “mouse click” procedure carried out by the authors within the R software, following the same concern shown in major point 3. Please, be careful with use of parameter general values in the QTL analysis. They might not explain the reality of current data.

In any case, there are three issues that I would like to highlight regarding this major point:

First, why would the selection of three markers as cofactors have necessarily been sufficient to control putative QTL outside the current interval or marker for any of the mapped populations? Would the inclusion of only three cofactors in the model have been sufficient to reduce residual variance and avoid any interference of this in the detected QTL and their main effects? Of course, cofactors should not be included in the model without any restriction, because the sample size would need to be sufficient to enable the estimation of all parameters. However, it is hard to accept that only three cofactors would have necessarily be sufficient to control “noise” from any of the situations.

Second, why was a window size of 10 centiMorgan (cM) used for CIM? Was this window size selected in the qtl R package considering the choice of default parameters? Would this window size have made sense for the rice experimental population (real data) considering the patterns of linkage disequilibrium (LD) for this crop species? Authors should mainly keep in mind the latter to perform QTL analysis. Maybe different window sizes would need to have been tested to evaluate the most consistent results.

Third, I honestly believe that CIM would not have actually been necessary in the present study. The authors have mentioned that CIM was used for detecting QTL main effects. However, why didn’t they consider such QTL main effects from MIM, since there was a great motivation to compare SHAP-XGB with the latter due to its estimation of complex interactions (epistasis) between QTL? Sincerely, it is hard to understand this authors’ strategy in the manuscript. Please, address this very relevant point from the study;

6)

MIM was performed using the QTL Cartographer software. In the subsection “Multiple interval mapping”, authors mentioned that the current software terminated abnormally in nine of the 40 established replications, being 10 replications used for each of the four simulated scenarios. In this context, there are four important issues from my point of view: (i) what would it explain this abnormal ending of MIM within the QTL Cartographer? Were the nine terminated replications related to different simulated scenarios or to a unique simulated scenario?; (ii) why did the authors only run MIM for four of the six simulated scenarios?; (iii) was the number of replications for each scenario really sufficient to obtain reliable means for the estimated parameters (accuracy of QTL detection) and suitable variation for a comparison with other approaches?; and (iv) where were the results from MIM really presented in the section “Results”? I didn’t find them in any figure of the manuscript, with the aim of comparing this approach with other methods and mainly with SHAP-XGB. Honestly, I didn’t understand the comparison procedure adopted by the authors. Wouldn’t the comparison mainly between MIM and SHAP-XGB have been the authors’ main motivation in the present study, considering that the former represents the main approach of QTL mapping which takes into account the interaction effects (epistatis) between QTL? It is totally unclear for readers. Without a clear comparison involving both approaches, it would be very hard to understand which was the real authors' motivation following their ideas from the introductory section;

7)

Considering the BayesC-based QTL mapping, authors have selected the default values for the hyperparameters of the model within the VIGoR R package. Why did the authors proceed like that? A consistent explanation should be provided to readers in the manuscript, so that they could not interpret the current selection as being just a "mouse click" procedure even within the R software;

8)

The last major point is related to the number of considered individuals for the analyzed populations. This is a very critical point in QTL mapping studies mainly when we are intended to estimate the effects of complex interactions (epistasis effects) between QTL. Although the authors have focused their study on rice heading date, which is an example of trait whose epistatic interactions have been elucidated, it is hard to believe that an experimental population of 174 individuals (and 200 for simulated populations) would really be helpful for the purpose of the present study. Using MIM, it is often challenging to detect statistically significant interactions between QTL based on a small sample size and considering a multiple dimensional genome search, which would reduce statistical power. Therefore, from my point of view, authors should rely their study on populations consisting of more than 300 or 400 individuals, which could provide an appropriate comparison among the current approaches, mainly between MIM and SHAP-XGB;

MINOR POINTS:

1) Lines 18, 19, and 32: try to use "QTL" instead of "QTLs". Please, check for this term throughout the entire manuscript;

2) Line 36: Association mapping? Isn't the current study based on QTL mapping and controlled experimental populations derived from biparental crosses?;

3) Lines 43-47: the first two sentences from the section "Introduction" need to be improved for readers. What would it be "efficient selection based on QTL genotypes"? Why did the authors refer to "development of medical technologies" if their current study is based on plants? What would these "black boxes" be in the context of quantitative genetics? Please, make clear these sentences in the manuscript;

4) Line 59: MIM was first proposed by Kao et al. (1999). Of course, authors can reference the paper of Laurie et al. (2014), which is a very relevant study in the context of MIM and has been effectively explored by the authors in the current work. However, the citation of Kao et al. (1999) is necessary. Please, include it in the manuscript;

5) Line 63: use "GWAS" instead of "GWASs". The former is used for both singular and plural;

6) Lines 63-64: authors provide a comparison between GWAS and QTL mapping based on inbred line crosses. That's alright. However, just a reminder: QTL mapping has already been proposed and extended in the context of an F1 segregating population, which can be obtained from the cross between heterozygous parents. If authors are interested, see the paper of Gazaffi et al. (2014). They propose an extension of the

traditional CIM for an F1 segregating population (full-sib progeny) and could be helpful in future studies.

### Decision letter (for ver. 2)

### Theoretical and Applied Genetics

Dear Prof. Onogi,

We have received the reports from our advisors on your manuscript TAAG-D-24-00529 "Applying gradient tree boosting to QTL mapping with Shapley additive explanations".

With regret, I must inform you that, based on the advice received, I have decided that your manuscript cannot be accepted for publication in Theoretical and Applied Genetics.

Below, please find the comments for your perusal. You are kindly requested to also check the website for possible reviewer attachment(s).

I would like to thank you very much for forwarding your manuscript to us for consideration and wish you every success in finding an alternative place of publication.

With kind regards,
Jiankang Wang
Editor
Theoretical and Applied Genetics

### COMMENTS TO THE AUTHOR:

The authors made great efforts revising their manuscript, based on the editor and reviewers' comments from previous review. However, I am sorry to say there are still some unsolved issues, which make it less easy to see the significant contribution of this study to QTL mapping theory and methods. I am not an expert on machine learning. Therefore, I cannot judge the suitability of applying the gradient tree boosting in QTL mapping. My evaluation is purely from the genetics / quantitative genetics point of view.

1. From Introduction, it seems the new method would contribute to the mapping of epistatic QTLs, and majority of this study would focus on epistasis, i.e. interaction between genes at different loci. But there are significant texts and contents on the mapping of additive (and dominant) QTLs.

2. MIM identifies the interactions between those QTLs with additive (and dominant) effects. In reality, epistasis can occur between the QTLs each having no additive (and dominant) effects. MIM has not been frequently used in QTM mapping studies. In fact, comparison should be made with most popular methods and software in QTL mapping. As an example, epistatic QTL mapping can be conducted in QTL IciMapping, an integrated software package which has been more widely used (Zhang et al., 2008, 2012; Li et al., 2008; Meng et al., 2015). QTL detection power and false discovery rate (FDR) can also be evaluated in the package as well from simulated mapping populations by defining a number of QTLs on the trait of interest.

3. One major disadvantage of IM is the ghost QTL phenomena at the presence of linkage. The one QTL on each chromosome model as has been used in simulation studies needs some justification. For simple models, the difference from different mapping methods may not be observed.

4. In simulation studies, it is not clear whether the additive (and dominant) effects are the same for all QTLs; whether the epistatic effects are the same for all interacting QTLs.

5 In simulation studies, each predefined QTL should have a detection power; each pair of interacting QTLs should have a detection power (Li et al., 2010; Zhang et al., 2012). It is not clear what the positive rates were referred to in Fig. 2 and Fig. 3. In addition, the estimation of positions and effects are also important in QTL mapping. I failed to see any results on this aspect.

6 TAG is not a journal in the field of physics or mathematics. In some formulas, well-accepted symbols should be adopted, or the expression and format need improvement. E.g., L139: Does |S| stand for the size of set S, |S|! stand for the factorial number of the size of set S? L149, does N¥{i} stand for the subset of N exclude element i?

Zhang, L., H. Li, Z. Li, and J. Wang. 2008. Interactions between markers can be caused by the dominance effect of QTL. Genetics 180: 1177-1190.

Li, H., J.-M. Ribaut, Z. Li, and J. Wang. 2008. Inclusive composite interval mapping (ICIM) for digenic epistasis of quantitative traits in biparental populations. Theor. Appl. Genet. 116: 243-260.

Li, H., S. Hearne, M. Bänziger, Z. Li, and J. Wang. 2010. Statistical properties of QTL linkage mapping in biparental genetic populations. Heredity 105: 257-267.

Zhang, L., H. Li, J. Wang. 2012. Statistical power of inclusive composite interval mapping in detecting

digenic epistasis showing common F2 segregation ratios. J Integr. Plant Biol. 54: 270-279
Meng, L., H. Li, L. Zhang, J. Wang. 2015. QTL IciMapping: Integrated software for genetic linkage map construction and quantitative trait locus mapping in bi-parental populations. Crop J. 3: 269-283

### Decision letter (for ver. 3)

### G3: Genes|Genomes|Genetics

Dear Dr. Onogi:

Thank you for submitting your manuscript entitled "Applying gradient tree boosting to QTL mapping with Shapley additive explanations" for peer review at G3: Genes|Genomes|Genetics, a fully open access journal published by the Genetics Society of America. We appreciate your interest in the journal.

Members of our Editorial Board have evaluated your manuscript and have determined that it is unlikely to fare well in peer review in its current form.

We downloaded everything and tried to run the R code. You have not set up the installation of all the dependencies correctly- so unless the user has all the packages of SHAPforxgboost already installed the installation will fail. the R script provided is not a re-usable tool for others but the analysis script used for a re-analysis of some "sample data".

What is unclear is who is the target audience? What is the usefulness ? Is this a tool intended for others to use in their QTL work? If so, it needs to be packaged like a tool, in a standalone package for R, and then distributed as an R tool. The current script is a script that uses a different package for a single data analysis not a tool/package that can be re-used by others.

Also missing are biological insights gained from this analysis that led to a new discovery? What follow-up was made possible by this analysis? why would this be useful? What evidence is there that there is a benefit or gain from these observations? what difference will this make in practice?

The data used are several years old, is there follow-up data that demonstrate that the novel predictions of interactions are real?

if you are able to address all of these concerns, we may send a revised manuscript out for peer review- but it would need to be substantially different from this manuscript.

If you choose to revise and resubmit, please:

1. Follow this link to submit the revised manuscript: https://g3.msubmit.net/cgi-bin/main.plex?el=A3NQ4Gcq1A2Eei5I3A9ftd1ruGLHHoeO9xncFGtw2MAZ
2. Include a description of your revisions in the cover letter of your revised manuscript.
We hope that our above comments are valuable to you and please contact us if you have any questions.

Sincerely,

Dr. Lauren McIntyre
Editor in Chief
G3

### Decision letter (for ver. 4)

### Bioinformatics

Dear Prof. Onogi,

Thank you for submitting your article to Bioinformatics.

The reviews of your manuscript are now in hand. The reviewers had substantial concerns about the manuscript and the Associate Editor, Russell Schwartz, has decided to reject your manuscript based on their advice.

The reviewers see your manuscript as rigorous and well written, with potential for your methods to be a useful tool for the community. However, they also raise some notable concerns. Some of these appear to be easily addressable questions of clarifying the contribution, but others are more fundamental concerns about significance and innovation with your paper seen as making an incremental advance on some related prior art. These are difficult issues to resolve in revision, which leads me to believe the paper will not meet the journal's high standards on these points even after a major revision.

As the journal receives more publishable manuscripts than its space will accommodate acceptance must be limited to manuscripts receiving the most favourable recommendations from reviewers.

Unfortunately, I must decline your request to have the manuscript published in Bioinformatics.

THE REVIEW COMMENTS can be found at the foot of this e-mail. We will not consider any further versions of this manuscript, but I hope that you find the comments useful should you decide to revise your manuscript for another journal.

Although we think your manuscript may not be a good fit for Bioinformatics, we would encourage you to consider accepting a transfer offer to Bioinformatics Advances, which is a broad-scope, open-access journal dealing with multiple aspects of computational biology. Papers are judged on the basis of scope, purposefulness of the contribution and soundness of method and validation. The Editors-in-Chief of Bioinformatics Advances, Alex Bateman (EMBL-EBI) and Thomas Lengauer (Max Planck Institute for Informatics, Saarbrücken, Germany), encourage you to consider this option. To learn more about Bioinformatics Advances, please visit https://academic.oup.com/bioinformaticsadvances
The transfer option is fast as the editors of Bioinformatics Advances are often able to use the reviewers' comments to arrive at a fast editorial decision. If there are at least two sufficiently good quality reviews included with the transfer, the editors of Bioinformatics Advances may be able to make an immediate decision regarding acceptance.

If you choose to transfer your article, the manuscript and any reviews will be sent directly to the editorial office of Bioinformatics Advances and you will receive a confirmation email. Bioinformatics Advances aims to provide a decision on transferred manuscript that have already been reviewed within three working days, and the journal's average time to first decision is 30 days. If your manuscript is suitable for the journal you will be provided details about any revisions required by the Bioinformatics Advances Editor based on the existing reviews. You will have an opportunity to revise your manuscript after transfer and before any further peer review.

Please note that Bioinformatics Advances is editorially independent of Bioinformatics and therefore the offer to transfer your manuscript does not imply that it will necessarily be sent for peer review or accepted by Bioinformatics Advances. However, the acceptance rate for transferred manuscript is favorable. If your article is accepted for publication in Bioinformatics Advances, an article processing charge will apply. Details can be found in the Instructions for Authors: https://academic.oup.com/bioinformaticsadvances/pages/instructions-to-authors

If you click below to approve, your paper will be moved to Bioinformatics Advances automatically. IMPORTANT: Please ensure that all authors agree to transfer or not transfer before clicking. This transfer offer will expire in 60 days.

Approve transfer: *** PLEASE NOTE: This is a two-step process. After clicking on the link, you will be directed to a webpage to confirm. ***
https://mc.manuscriptcentral.com/bioinformatics?URL_MASK=56af6063dd85412da8b5ddfe001f16b9

Decline transfer: *** PLEASE NOTE: This is a two-step process. After clicking on the link, you will be directed to a webpage to confirm. ***

https://mc.manuscriptcentral.com/bioinformatics?URL_MASK=34b9c12a04544bb39f3336d713bec411

On behalf of the Executive Editor, I would like to thank you for considering Bioinformatics to present your work and I look forward to the possibility of receiving other manuscripts from you in the future. However, we will not be able to reconsider this manuscript again for the journal.

Best regards,
Alison Hutchins
Bioinformatics Editorial office

Here are the comments of the reviewers:
---------------------------------------

#### Reviewer: 1

Paper is well written and it shows nicely the added value why use of Shapley values may be a good idea. Paper may be nice add to the toolbox for biologists and breeders.
However, novelty of this work is not huge (cf. almost same topic of Johnsen et al. 2021). I have following quite general comments:

1) Different kind of tree models (CART, Random forest and so on) have been around for long time. You should add some more tree model-based references to your paper, which also indirectly indicate that novelty of the paper is not extremely high. Some suggested references:

McKinney BA, Reif DM, Ritchie MD, Moore JH. Machine learning for detecting gene-gene interactions: a review. Appl Bioinformatics. 2006;5(2):77-88. doi: 10.2165/00822942-200605020-00002.

Meijsen JJ, Rammos A, Campbell A, Hayward C, Porteous DJ, Deary IJ, Marioni RE, Nicodemus KK. Using tree-based methods for detection of gene-gene interactions in the presence of a polygenic signal: simulation study with application to educational attainment in the Generation Scotland Cohort Study. Bioinformatics. 2019 Jan 15;35(2):181-188. doi: 10.1093/bioinformatics/bty462.

Waldmann, P. Genome-wide prediction using Bayesian additive regression trees. Genet Sel Evol 48, 42 (2016). https://doi.org/10.1186/s12711-016-0219-8

Stephan, J., Stegle, O. & Beyer, A. A random forest approach to capture genetic effects in the presence of population structure. Nat Commun 6, 7432 (2015). https://doi.org/10.1038/ncomms8432

Montesinos López, O.A., Montesinos López, A., Crossa, J. (2022). Random Forest for Genomic Prediction.

In: Multivariate Statistical Machine Learning Methods for Genomic Prediction. Springer, Cham. https://doi.org/10.1007/978-3-030-89010-0_15

2) Throughout in your paper, please clarify when you discuss about pairwise epistasis (interaction of two QTLs) or higher-order epistasis (interaction between more than two QTLs). Currently you speak about epistasis without making any difference between these which difficults reading.

3) In your paper, QTL mapping and association mapping (GWAS) are presented as two separate problems (I agree that they can be so). However, for biparental crosses, If marker map is dense enough, they are very similar kind of problems because there is no need to go between markers (QTL interval mapping) anymore. Please clarify this aspect more in your paper. Alternatively, if you do not agree, please explain better why it is not so.

4) It may be unclear for the reader, why you can take method for genomic prediction (BayesC) and use it directly for QTL mapping as you did here. Please explain this more to the reader.

5) You mention about sure independence screening as filter to make your method more scalable. This is true, but another aspect of it is that it may be more powerful to do epistasis search in two phases by first finding main effects and then interactions. See e.g.,

Kärkkäinen HP, Li Z, Sillanpää MJ. An Efficient Genome-Wide Multilocus Epistasis Search. Genetics. 2015 Nov;201(3):865-70. doi: 10.1534/genetics.115.182444.

Mathew B, Léon J, Sannemann W, Sillanpää MJ, Detection of Epistasis for Flowering Time Using Bayesian Multilocus Estimation in a Barley MAGIC Population, Genetics, Volume 208, Issue 2, 1 February 2018, Pages 525–536, https://doi.org/10.1534/genetics.117.300546

#### Reviewer: 2

One main difference between QTL mapping and GWAS is whether one position between two linked markers can be identified as QTL/QTN. If YES, it is QTL mapping. In this manuscript about QTL mapping, please add the related theory and method to identify the QTLs between two linked markers. In Monte Carlo simulation studies, some QTLs should be located in the interval of two linked markers, while in real data analysis some significant QTLs and their true genes can be found in the interval of two linked markers.

In rice real data analysis, it is necessary to list the gene-by-gene interactions (GGI) identified in this manuscript and supported by some pieces of evidence in molecular biological experiments, in particular, new GGIs.

Lines 56-57: For the sentence of “Bayesian regression where whole genome-wide markers are used as explanatory variables simultaneously (Zhang and Xu 2005; Li and Sillanpää 2012)”, two references, Xu (2003) Genetics 163: 789–801 and Wang et al. (2005) Genetics 170: 465–480 , should be cited.

Lines 59-61: For the sentence of “Detection of QTL/gene interactions has been of interest in genome-wide association studies (GWAS), although detection is more difficult than in inbred line crosses because of the generally low allele frequencies”, one important reference, Li et al. (2022) Mol. Plant. 15(4), 630–650, should be cited.

In the Introduction section, the authors should focus on the methodological advances of QTL mapping rather than GWAS.

In Table 1, only additive effects and their epistatic effects were simulated. It is too simple. In F2 populations, additive and dominant effects and their interaction effects (aa, ad, da and dd) should be simulated.

In Figure, the AUC values of the y axis were more than 100%, which are not correct.

---------------------------------------

### Decision letter (for ver. 5)

### Breeding Science

Dear Onogi,

Reviewers have now commented on your paper. You will see that they are advising that you revise your manuscript. If you are prepared to undertake the work required, I would be pleased to reconsider my decision.

For your guidance, reviewers' comments are appended below.

Reviewers' comments:

### Editor:

This manuscript has been peer-reviewed by two reviewers specializing in the relevant field. As a result of the review, both reviewers have pointed out issues that require consideration. Please carefully address the reviewers' comments, perform additional analyses as necessary, and revise the manuscript accordingly. I would also like to provide the following comments for your consideration in conjunction with the reviewers' feedback:

The primary goal of QTL analysis is not only the detection of QTLs but also the estimation of their precise positions and effects. For the QTLs detected by the proposed method, it is necessary to examine how their accurate positions and effects can be estimated. If it is challenging to conduct a detailed analysis by applying specific methods to the data, this issue should be addressed as an unresolved challenge in the Discussion section.

In this study, each QTL is independently located on a separate locus. Therefore, main effects and interaction effects, such as epistasis, are likely to be independent in most cases. On the other hand, one of the main purposes of performing CIM or MIM is to determine whether multiple linked QTLs can be detected separately. In such cases, it is necessary to examine whether issues like the detection of a ghost QTL between two QTLs arise.

Finally, one of the advantages of machine learning approaches like XGBoost is their ability to capture complex interactions. In this study, interactions between two QTLs were considered, and genotypic values were simulated as their linear combination. However, for genes involved in metabolic pathways, more extensive and complex interactions may arise, such as upstream-downstream relationships within the same pathway or relationships mediated by alternative pathways. These are precisely the situations where machine learning methods like XGBoost might be well-suited. While it may be challenging to perform complex simulation analyses, it seems plausible that significant differences between XGBoost and linear statistical models in QTL analysis might emerge under such conditions.

#### Reviewer #1:

In this manuscript, the authors applied a popular and highly evaluated machine learning method, XGboost, to QTL analysis, where marker effects were estimated as importance of features in XGboost model assessed by SHAP, a Shapley value like score, and compared the method (SHAP-XGB) with conventional QTL methods including CIM, MIM, ICIM and BayesC for the efficiency in detecting QTL main and interaction effects. I think that it is important to evaluate the applicability of AI models such as gradient boosting and deep neural network to conventional statistical genetic analyses including QTL analysis and GWAS, thus, this study was of some interest to me. But there were some problematic parts in the manuscript. The authors should address the issues pointed out, as listed below.

Major points

Simulations section in Materials and Methods

1. Marker positions were not described. The authors stated that each chromosome with length 120cM included 20 markers but did not describe how markers were located. Were markers located equidistantly on

each chromosome?

2. Line 285: What did "markers located within 20cM of the QTL" mean? Was it the region of 20cM length including a QTL at midpoint? How many markers were included in the region?

3. I think that the numbers of true positive markers and true negative markers (not linked to QTL) were critical to drawing ROC curve in detecting QTL main effect and that the numbers of true positive marker pairs and true negative marker pairs (not linked to epistatic QTL pairs) were also critical for ROC curve in detecting QTL interaction effect. Please show the numbers of the true positive markers and the true positive marker pairs in a total of 160 markers and a total of 12720 (=160*159/2) marker pairs although the numbers of true positives were changed depending on simulation replication. Anyway, the authors are required to clearly describe how true and false positive rates were calculated.

4. Interval mapping methods, CIM, MIM and ICIM, can detect QTL not only at the marker position but on any position between markers. Were only marker positions tested for QTL with interval mapping methods in this study?

Mapping dominance effects section in Materials and Methods

5. Line 302: The authors stated that genotypes were differently corded according to the mode of gene action (additive, complete dominance and overdominance) for BayesC and SHAP-XGB. In this way, genome-wide QTL analysis is repeatedly performed for the same dataset by changing the genotype code according to the mode of gene action of QTL as the mode is actually unknown. It seems to me that this way of QTL analysis is much ineffective. Instead, it would be desirable that genotypes of QTL are separately coded for additive and dominance effect, i.e., (1,0,-1) for additive effect and (0,1,0) for dominance effect, and both additive and dominance effects are considered in the model of BayesC and SHAP-XGB as in CIM and MIM. But I don't know whether such implementation is possible in the available R version of BayesC and SHAP-XGB. Moreover, three types of QTL interactions, additive x additive, additive x dominance and dominance x dominance, could be incorporated in the model of BayesC and SHAP-XGB although the model dimension becomes much huge.

Minor points

1. Line 123: "By approximating f" should be written as "By approximating l".
2. Line 188: Calculation of SHAP interaction values should be included in Step 3.

#### Reviewer #2:

This paper explores the potential benefits of using gradient tree boosting with Shapley additive explanations (SHAP) to detect additive, dominance, and epistatic QTLs. The authors applied XGBoost, a gradient tree boosting method, to biparental populations and derived feature importance using the SHAP approach. They compared their proposed SHAP-XGB model to conventional QTL mapping methods, such as ICIM and Bayesian models, using both simulated and real rice datasets. Results showed that SHAP-XGB performed comparably to conventional models in detecting QTLs for additive and dominance effects and outperformed them in mapping QTL interaction effects. This suggests that the paper presents a valuable idea for efficiently detecting QTLs across various gene action types. The authors also provide a thoughtful discussion of SHAP-XGB's potential extensions and limitations, including its application to GWAS. While the potential extensions and limitations are well discussed, some further discussion regarding simulation settings is needed.

I attached a PDF file, so please refer to the file for detailed comments.

=PDF file=

**Comments on: Onogi et al., 2024, Applying gradient tree boosting to QTL mapping with Shapley additive explanations, submitted for Breeding Science.**

**General Comments:**

This paper explores the potential benefits of using gradient tree boosting with Shapley additive explanations (SHAP) to detect additive, dominance, and epistatic QTLs. The authors applied XGBoost, a gradient tree boosting method, to biparental populations and derived feature importance using the SHAP approach. They compared their proposed SHAP-XGB model to conventional QTL mapping methods, such as ICIM and Bayesian models, using both simulated and real rice datasets. Results showed that SHAP-XGB performed comparably to conventional models in detecting QTLs for additive and dominance effects and outperformed them in mapping QTL interaction effects. This suggests that the paper presents a valuable idea for efficiently detecting QTLs across various gene action types. The authors also provide a thoughtful discussion of SHAP-XGB's potential extensions and limitations, including its application to GWAS.

In the following, I offer brief comments on areas that may help improve the manuscript.

**Short Comments:**

**P7, L176**

“i” -> “*i*” (please make it italic)

**P8, L180**

Please provide citation for the R package “xgboost”.

**P11, L259 - 263**

Please provide more detailed information on how to generate marker genotype data for F2 and RIL populations.

**P11, L265 - 266**

How was the ratio of variance between additive and epistatic effects determined? The actual magnitude of variance is shown in the subsequent explanations and tables for each scenario, but please provide information on how you initially set this ratio.

**P11, L276 - 278; P12, L302 - 303; P17, L419 - 423**

The study provided scenarios assuming only additive or dominance (and their interaction) effects. However, as mentioned in the Discussion, SHAP-XGB targeting additive effects can map dominance effects or dominance-by-dominance interaction effects. This may hinder accurate identification of gene actions for the detected QTLs. I suggest including an additional scenario that assumes both additive and dominance effects to evaluate how SHAP-XGB performs in detecting these effects simultaneously. For this scenario, proper genotype coding is crucial. It may be beneficial to incorporate two or three types of genotype coding as inputs for XGBoost concurrently.

**P15, L370 - 371**

"Supplemental Fig. 5" -> "Supplemental Fig. 4" ?

**Others**

Although not addressed in this study, it would be interesting to explore SHAP-XGB's performance on binary traits. The nature of gradient tree boosting suggests that this method could be readily adapted to classification problems, potentially allowing for more intuitive detection of QTLs—including those with epistatic effects—associated with binary traits than current methods. Incorporating such scenarios and discussions could significantly enhance the study's impact and applicability.

**Conclusion:**

Detecting QTLs with additive, dominance, and epistatic effects is crucial in genetics as it can unravel the genetic architecture of complex traits. This paper deserves praise for applying a machine learning method to achieve this goal by introducing an interpretable approach. The proposed model outperformed conventional models and achieved desirable results in both simulated and real datasets. While the potential extensions and limitations are well discussed, some further discussion regarding simulation settings is needed.
